# Born with intronless ERF transcriptional factors: C_4_ photosynthesis inherits a legacy dating back 450 million years

**DOI:** 10.1101/2022.10.14.512192

**Authors:** Ming-Ju Amy Lyu, Huilong Du, Hongyan Yao, Zhiguo Zhang, Genyun Chen, Faming Chen, Yong-Yao Zhao, Qiming Tang, Fenfen Miao, Yanjie Wang, Yuhui Zhao, Hongwei Lu, Lu Fang, Qiang Gao, Yiying Qi, Qing Zhang, Jisen Zhang, Tao Yang, Xuean Cui, Chengzhi Liang, Tiegang Lu, Xin-Guang Zhu

**Affiliations:** State Key Laboratory of Plant Molecular Genetics, Center of Excellence for Molecular Plant Sciences, Chinese Academy of Sciences, Shanghai, China, 200032; State Key Laboratory of Plant Genomics, Institute of Genetics and Developmental Biology, Innovation Academy for Seed Design, Chinese Academy of Sciences, Beijing, China; University of Chinese Academy of Sciences, Beijing, China; School of Life Sciences, Institute of Life Sciences and Green Development, Hebei University, Baoding, China; State Key Laboratory of Genetic Engineering, School of Life Sciences, Fudan University, Shanghai 200438, China; Biotechnology Research Institute/National Key Facility for Gene Resources and Gene Improvement, Chinese Academy of Agricultural Sciences, Beijing, 100081, China; Center for Genomics and Biotechnology, Fujian Provincial Key Laboratory of Haixia Applied Plant Systems Biology, Key Laboratory of Sugarcane Biology and Genetic Breeding, National Engineering Research Center for Sugarcane, College of Life Sciences, Fujian Agriculture and Forestry University, Fuzhou, China; China National GeneBank, Shenzhen, 518120, China

**Keywords:** *Flaveria* genome, C_4_ photosynthesis, Tandem duplication, Intronless ERF TFs

## Abstract

The genus *Flaveria*, containing species at different evolutionary stages of the progression from C_3_ to C_4_ photosynthesis, is used as a model system to study the evolution of C_4_ photosynthesis. Here, we report chromosome-scale genome sequences for five *Flaveria* species, including C_3_, C_4_, and intermediate species. Our analyses revealed that both acquiring additional gene copies and recruiting ethylene responsive factor (ERF) *cis*-regulatory elements (CREs) contributed to the emergence of C_4_ photosynthesis. ERF transcriptional factors (TFs), especially intronless ERF TFs, were co-opted in dicotyledonous C_4_ species and monocotyledonous C_4_ species in parallel. These C_4_ species co-opted intronless ERF TFs originated from the Late Ordovician mass extinction that occurred ∼450 million years ago in coping with environmental stress. Therefore, this study demonstrated that intronless ERF TFs were acquired during the early evolution of plants and provided the molecular toolbox facilitating multiple subsequent independent evolutions of C_4_ photosynthesis.

## Introduction

C_4_ photosynthesis is a complex trait that evolved from ancestral C_3_ types in the last 35 million years (Sage, 2004; Sage et al., 2012). With high light, water, and nitrogen use efficiencies (Vogan and Sage, 2011; Zhu et al., 2008), C_4_ photosynthesis is an ideal target to be engineered into C_3_ crops to increase crop yield (Long et al., 2015; Maurino and Weber, 2013; Zhu et al., 2010). Compared to C_3_ photosynthesis, C_4_ photosynthesis dedicates more genes to carbon fixation, and these genes are compartmentalized either in mesophyll cells (MCs) or in bundle sheath cells (BSCs) (Hatch, 1987; Slack and Hatch, 1967). These MCs and BSCs form the specialized C_4_ “Kranz anatomy” (Hatch, 1987). Therefore, the evolution of C_4_ photosynthesis requires modifications of both metabolism and leaf anatomy in C_3_ ancestors. Though complex, C_4_ photosynthesis has evolved independently more than 70 times in angiosperms, making it an excellent example of convergent evolution of a complex trait (Sage, 2016). How such a complex trait emerges repeatedly remains an unresolved question in biological research.

All genes that function in C_4_ photosynthesis have counterparts in C_3_ species (Christin et al., 2013; Christin et al., 2009; Moreno-Villena et al., 2018; Williams et al., 2012). The same C_4_ orthologous genes that show relatively high transcript abundances were co-opted in different C_4_ lineages in parallel (Emms et al., 2016; Moreno-Villena et al., 2018). Moreover, the recruited C_4_ genes often adopt pre-existing regulatory mechanisms of photosynthesis (Burgess et al., 2016), which enables the coordinated expression of C_4_ cycle genes with other photosynthesis-related genes. This recruitment of pre-existing elements provides a mechanism for the repeated emergence of C_4_ photosynthesis in independent lineages. In available genome sequences, conserved *cis*- regulatory elements (CREs) as well as transcription factors (TFs) have been identified that control the MCs or BSCs specificity of C_4_ genes in different monocotyledonous C_4_ lineages (Burgess et al., 2019; Gupta et al., 2020; John et al., 2014). Moreover, conserved TFs controlling MCs and BSCs specificity of gene expression were also identified between monocotyledonous and dicotyledonous C_4_ species (Aubry et al., 2014). These observations nevertheless raise the questions of when and how these shared regulatory mechanisms were co-opted into the different C_4_ species that diverged 160 million years ago (mya) (Kumar et al., 2017), which is much earlier than the emergence of C_4_ photosynthesis, *i.e.*, 35 mya (Sage et al., 2011).

Among the dicotyledonous model systems for C_4_ photosynthesis, the genus *Flaveria* is remarkable because it contains C_3_, C_4_, and many intermediate species (Powell, 1978). During the last decades, studies based on this genus have contributed to our current understanding of the evolution of C_4_ photosynthesis (Gowik and Westhoff, 2011; Powell, 1978; Sage et al., 2013). However, due to a lack of genome reference, current knowledge of the regulation of photosynthesis genes in this genus is still very limited. The first version of *Flaveria* genome references, including four species, were published recently (Taniguchi et al., 2021), and provide valuable resources for protein coding gene predictions for this genus. However, as these genomes were generated using short-read whole genome-sequencing, the assembled genomes are fragmented, which compromises their potential application (Taniguchi et al., 2021). Taking advantage of long-read genome sequencing technology, here we reported chromosome-scale genome references of five *Flaveria* species, with which we conducted a systematic study of CREs and TFs during the evolution of C_4_ photosynthesis. We found that ethylene responsive factor (ERF) CREs were recruited by C_4_ photosynthesis during evolution. Moreover, intronless ERF TFs that originated 450 mya to cope with environmental stress were recruited into different C_4_ lineages; furthermore, our study highlighted the role of intronless ERF TFs in the evolution and regulation of C_4_ photosynthesis, and provided a mechanism underlying the repeated evolution of C_4_ photosynthesis.

## Results

### Analysis of five *Flaveria* genome assemblies showed that transposable elements were more abundant in the C_4_ species than other species

The genome sequences of five *Flaveria* species, *i.e., F. robusta* (Frob, C_3_), *F. sonorensis* (Fson, C_3_-C_4_), *F. linearis* (Flin, C_3_-C_4_), *F. ramosissima* (Fram, C_3_-C_4_) and *F. trinervia* (Ftri, C_4_) were obtained with PacBio RSII single-molecule real-time (SMRT) sequencing technology (Figure 1a). The assembled genome size was gradually increased during the evolution of C_4_ photosynthesis in this genus, from 0.55 Gb in the C_3_ species Frob, to 1.26∼1.66 in the C_3_-C_4_ species, and to 1.8 Gb in the C_4_ species Ftri (Table S1), and these data were consistent with the analysis based on flow cytometry (Supplemental Note 1). Based on chromatin conformation capture (Hi-C seq), 98% to 99% of the assembled genome sequences were anchored to 18 pseudo-chromosomes (Figure S1 and Supplemental Note 2), which was supported by fluorescence *in situ* hybridization (FISH) results in Frob, Flin, and Ftri (Figure 1a). This was consistent with the reported chromosome number of 36 (2n) in these five *Flaveria* species (Powell, 1978). Genome completeness was estimated using Benchmarking Universal Single-Copy Orthologues (BUSCO) genes and resulted in coverage from 92.5% to 99.2% of the BUSCO genes. Additionally, an average RNA-seq reads mapping rate of 94.3% (from 86.7 to 97.3%) was obtained (Supplemental Note 3), suggesting completeness and high quality of genome assemblies for the five *Flaveria* species.

**Figure 1.**
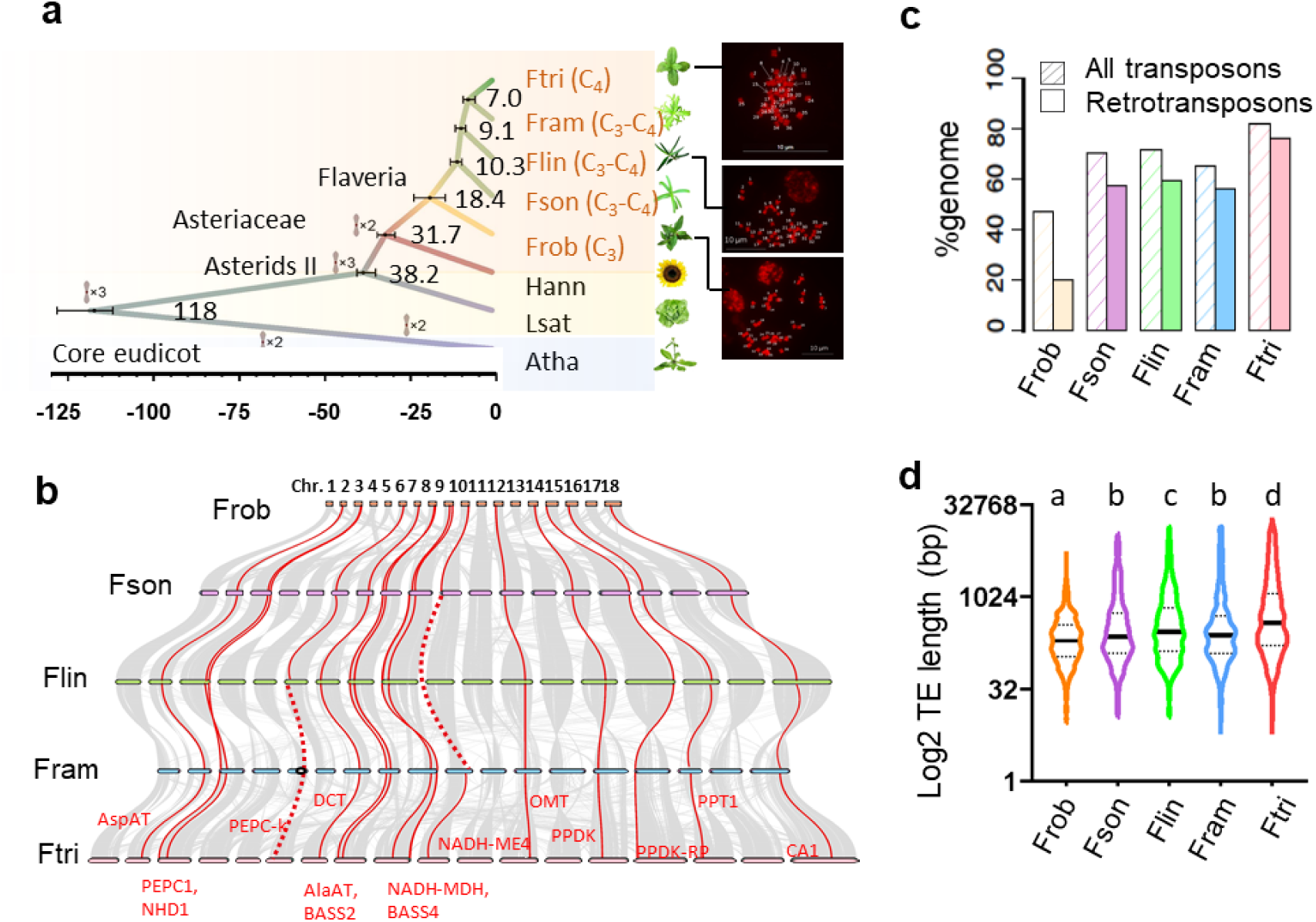
Transposon elements contributed to enlargement of genome size and promoters of C_4_ genes during *Flaveria* evolution. (a) Summary of phylogeny and timescale of the five *Flaveria* species and the three indicated outgroup species. Bars represent 95% confidence intervals of the estimated divergence time. Whole genome duplications are shown at the corresponding node/branch. Panels at the right display fluorescence *in situ* hybridization images to assess the chromosome numbers in Ftri, Flin, and Frob. (c) Collinearity of chromosomes among *Flaveria* species. C_4_ genes are drawn in red line. Dashed lines represent either failure in anchoring to chromosome (NADP-ME in Flin) or a deletion from the genome (PEPC-k in Fram). (b) Proportions of transposon elements, relative to the whole genome by length. (d) Assessment of 15 C_4_ genes (from panel c), showing that the C_4_ species Ftri has relatively longer TEs in the promoter region (3 kb upstream of start codon at the 5’ end) of these loci. (Abbreviations: Frob: *F. robusta*, Fson: *F. sonorensis*, Flin: *F. linearis*, Fram: *F. ramosissima*, Ftri: *F. trinervia*.)

Although genome size was tripled in the C_4_ species Ftri compared to the C_3_ species Frob, the number of protein coding genes was comparable between the C_3_ and C_4_ species, with 35,875 (Frob) and 32,915 (Ftri) respectively, and 37,028 to 38,652 protein coding genes were predicted in the C_3_-C_4_ species (Table S1). We compared the predicted protein coding genes from our assembly with those from Taniguchi’s assembly (Taniguchi et al., 2021), we found that around 96% protein coding genes were overlapped between our assembly and Taniguchi’s assembly (Taniguchi et al., 2021) (Supplemental Note 4). Therefore, the annotated protein-coding genes in this study can be considered reliable.

The chromosome-scale assembly of genome sequences and reliable gene annotations allowed us to study the evolution of known C_4_ enzymes and C_4_ transporters (termed as C_4_ genes) on location on the chromosomes. We identified eight enzymes and seven transporters as C_4_ versions by combining the gene phylogenetic tree and transcript abundances (Supplemental Note 5). As C_4_ versions of C_4_ genes, but not their orthologs, were reported to be induced by light in C_3_ species, we thus verified the C_4_ version of the C_4_ enzymes by examining their responsiveness to light. The C_4_ versions of C_4_ genes appeared quickly (after 2 hours) and were up-regulated after 4 hours in C_4_ species after being illuminated, and such light responses were intermediate in the C_3_-C_4_ species (Figure S2), which suggested the accuracy of identification of the C_4_ versions of C_4_ genes, and also revealed a gradual gain of light responsiveness during C_4_ evolution. The synteny of the 18 chromosomes was conserved in the five *Flaveria* species; from 50% to 75% of protein coding genes were colinear between Frob and the other species (Figure 1b and Supplemental Note 2). Notably, the chromosome locations of all 15 C_4_ version of C_4_ genes were conserved during evolution (Figure 1b).

Transposable elements (TEs) showed the highest abundance in the C_4_ species, where they accounted for 82% of the total genome, followed by C_3_-C_4_ species (from 65.6% to 71.8%), whereas that percentage in the C_3_ species was 47.1% (Figure1c and Supplemental Note 6). In all five species, long terminal repeat retrotransposons (LTR-RTs) comprised the majority of the TEs, accounting for an average 76% of the total TEs (from 42% to 91%) (Figure 1c). C_4_ genes had longer TEs on the promoter regions in the C_4_ species than their counterparts in C_3_ and C_3_-C_4_ species do (Figure 1d).

### C_4_ genes acquired elevated protein levels during evolution, which were regulated mainly at transcriptional levels

The transcript and protein abundances of C_4_ genes were generally higher in C_4_ species than in C_3_ and C_3_-C_4_ species (Figure 2a). To study whether transcriptional or translational regulation is primarily responsible for the observed difference in protein abundance between species with different photosynthetic types, we compared the protein-to-mRNA ratios (PTR) between genes in five *Flaveria* species. Low PTR genes and high PTR genes were defined as genes with PTR less than the mean PTR minus standard deviation (SD) and higher than the mean PTR plus SD, respectively (Figure 2b), and the remaining genes were defined as moderate PTR genes. In general, there was a positive correlation between mRNA and protein levels, with Pearson correlations ranging from 0.27 to 0.45, and most genes had moderate PTRs (Figure 2c). An average of 166 low PTR genes (from 121 to 201) and 395 high PTR genes (from 375 to 462) were obtained in the five species (Supplemental Notes 7∼9). In the C_4_ species, seven C_4_ genes were characterized as low PTR genes, whereas three or fewer C_4_ orthologous genes were low PTR genes in the C_3_ and C_3_-C_4_ species.

**Figure 2.**
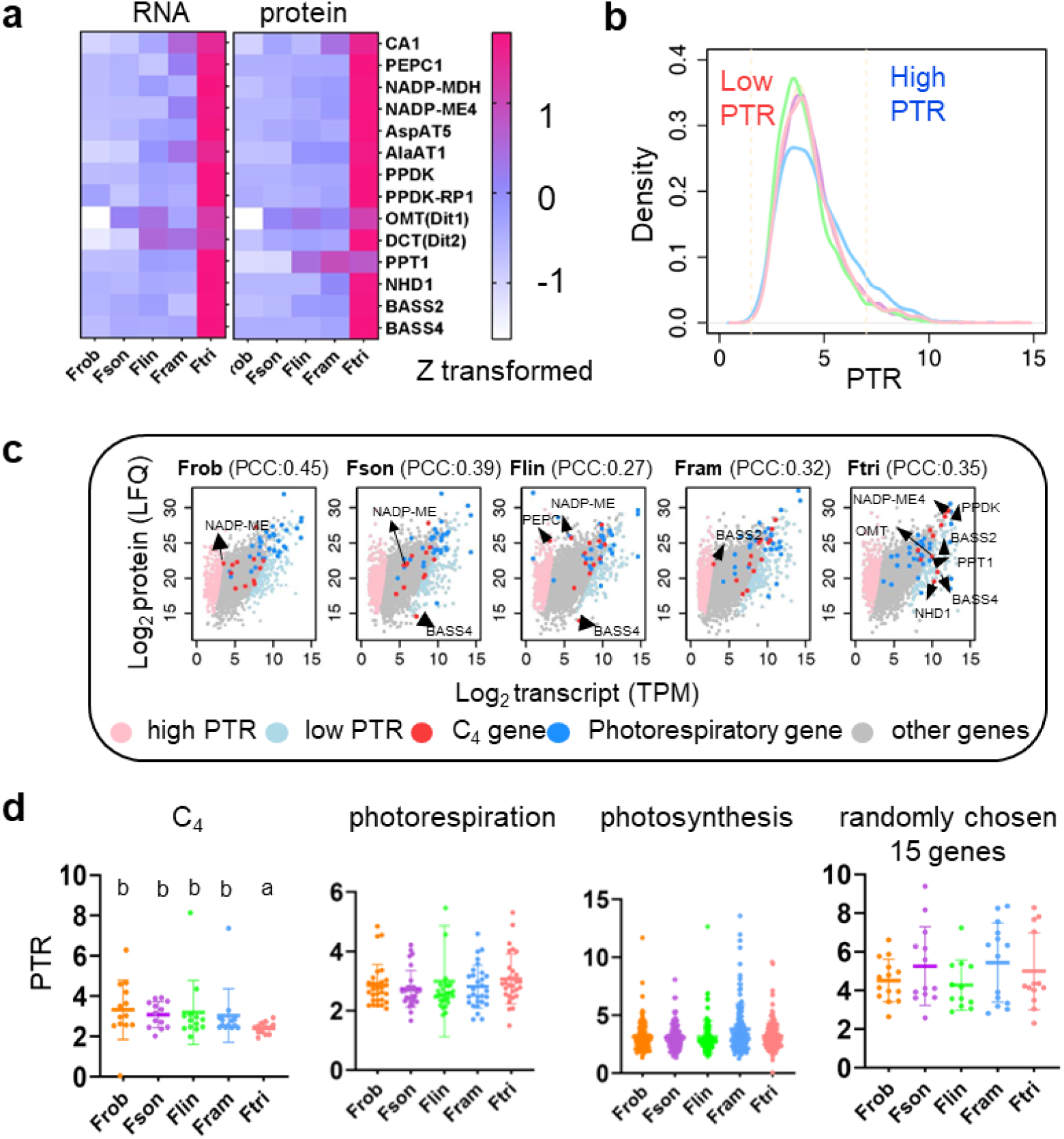
The C_4_ species had increased transcript abundances of C_4_ genes. (a) RNA-seq and proteomics data for the C_4_ genes in the five *Flaveria* species show increased transcript and protein abundances of C_4_ genes in the C_4_ species Ftri. (b) The protein-to-mRNA ratio (PTR) distribution of genes from the five *Flaveria* species. High PTR and low PTR genes are defined as genes with PTR higher than the mean plus one standard deviation (SD) and with PTR values lower than the mean minus one SD respectively. (c) Scatter plot of protein versus transcript abundance of the five *Flaveria* species. low PTR and high PRT C_4_ genes were labeled with arrows. Note the trend towards lower PTR for the C_4_ gene set in Ftri, as compared to the C_3_ Frob and the three intermediate *Flaveria* species. In contrast, there is no apparent shift in PTR for photorespiratory genes. (d) PTR values for the C_4_ gene set in the five *Flaveria* species, showing that C_4_ genes have significantly lower PTR in C_4_ species Ftri than in the C_3_ Frob or the three intermediate species. Note that no such decrease is showed for photorespiratory genes, photosynthesis genes and randomly chosen genes. (Abbreviations for proteins see Supplemental Note 8, Figure S14.)

The low PTR genes were enriched in gene ontology (GO) of photosynthesis and photosynthesis related GO terms, including chloroplast, PSII, and others (Supplemental Note 9), consistent with an early study in *Arabidopsis thaliana* (Atha), which showed that the photosynthesis related genes had significantly lower PTRs than other genes in photosynthetic functional leaf tissues (Mergner et al., 2020) (Supplemental Note 9). C_4_ genes showed significantly lower PTRs in the C_4_ species than their orthologs in the C_3_ and C_3_-C_4_ species did (Figure 2d), whereas photorespiratory genes or photosynthesis genes (not including C_4_ genes) showed comparable PTRs across the five *Flaveria* species. Therefore, during the evolution of C_4_ photosynthesis, C_4_ species acquired elevated protein levels for C_4_ genes, which were regulated mainly at transcriptional levels.

### Tandem duplication and recruitment of ERF *cis*-regulatory elements contributed to the increased transcript abundances of C_4_ genes

We then analyzed what contributed to the increased transcript abundance of C_4_ genes in the C_4_ species. Carbonic anhydrase (CA), phospho*enol*pyruvate carboxylase (PEPC), and PEPC kinase (PEPC-k) showed extra copies in C_4_ species, which were derived from tandem duplications (Figure 3a and Figure S3). For example, the C_4_ version of PEPC, termed PEPC1 because it showed the highest transcript abundance among the other paralogs in C_4_ species, had three copies in the C_4_ species Ftri, and only one copy in the other species. The three paralogs of PEPC1 in Ftri, termed as PEPC1.1, PEPC1.2, and PEPC1.3, were located on the same chromosome (Chr3) (Figure 3b). The existence of the three PEPC1 paralogs on the chromosome was further verified by PCR (Supplemental Note 10). In Ftri, all three PEPC1s had comparable transcript abundances, which were higher than those in the other four species (Figure 3b). Additionally, they were all upregulated under long-term low CO_2_ treatment (100 ppm) compared to normal CO_2_ conditions (380 ppm), suggesting that these triplets hosted shared regulatory mechanisms. Indeed, all three paralogs harbored the mesophyll expression module 1 (MEM1) CRE (Akyildiz et al., 2007) ( Supplemental Note 10); moreover, the three paralogs showed similar signatures of chromatin accessibility through transposase-accessible chromatin using sequencing (ATAC-seq) (Figure 3c and Supplemental Note 11).

**Figure 3.**
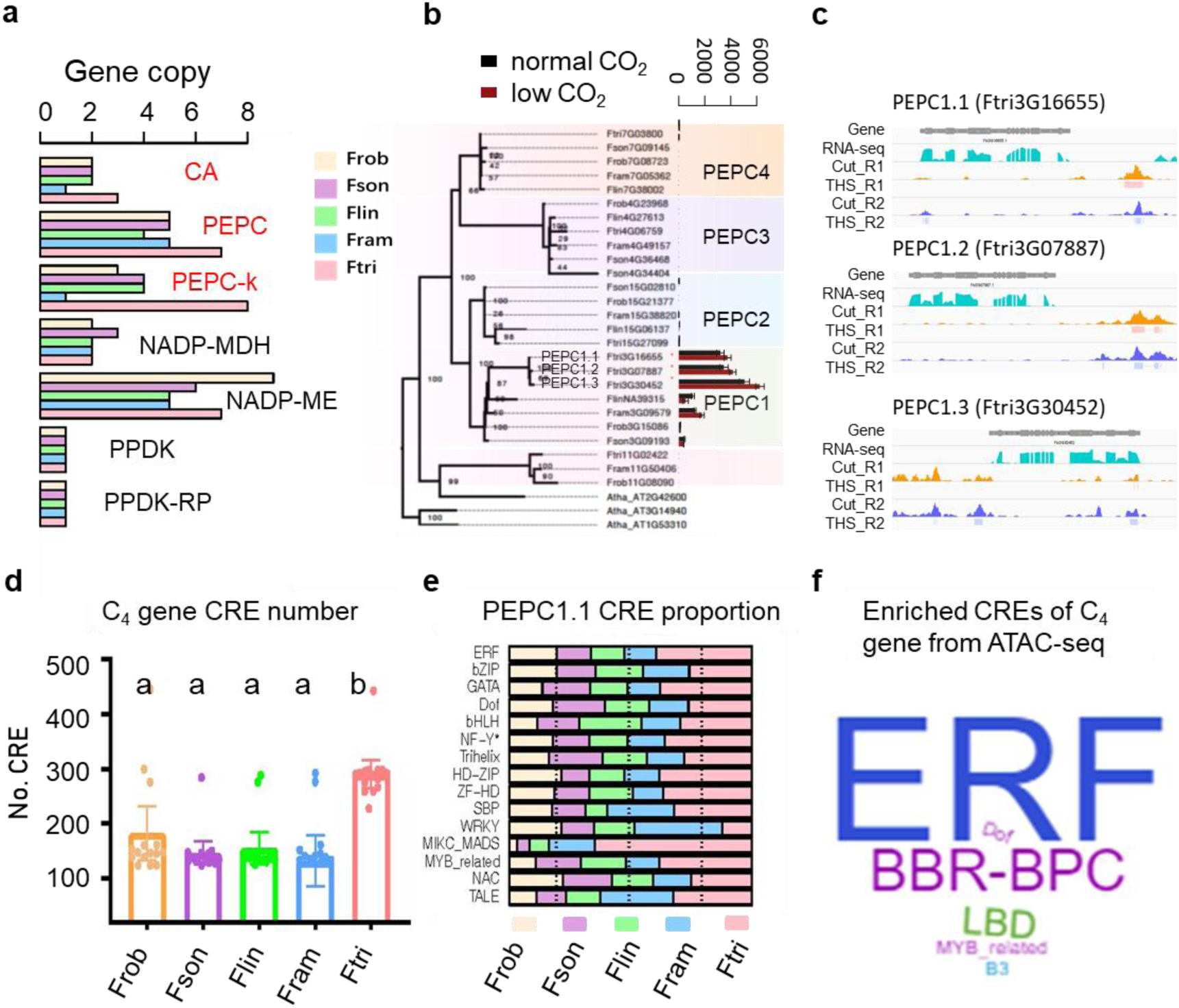
Tandem duplications and recruitments of ERF *cis*-regulatory elements contributed to the increased transcript abundances of C_4_ genes in the C_4_ species Ftri. (a) Copy number of the C_4_ version of C_4_ enzymes in five *Flaveria* species. Note that CA, PEPC, and PEPC-K have more copies in the C_4_ species Ftri than other *Flaveria* species. (b) Gene tree of PEPC orthologs, PEPCs from *Arabidopsis thaliana* (Atha) are used as outgroups. PEPCs in *Flaveria* species are categorized into four groups, and PEPC1 is the C_4_ version according to the highest expression levels among all PEPCs. PEPC1 has three copies in the C_4_ species Ftri, showing comparable transcript abundances in leaves and uniform upregulation when plants were grown under low CO_2_ conditions (100 ppm) compared to normal CO_2_ conditions (380 ppm). (c) Integrated Genome Viewer (IGV) of RNA-seq reads and ATAC-seq reads of three PEPC1 in Ftri. Tn5 cuts and transposase hypersensitive sites (THS) from two biological replicates show that the three PEPC1 have shared chromatin accessibility upstream of their coding region. (d) Bar plots show the number of predicted *cis*-regulatory elements (CREs) from the promoter region (3kb upstream of start codon) of all C_4_ genes in the five *Flaveria* species. (e) An example of the distribution of the top 15 CREs in C_4_ genes, noting that Ftri has more CREs in PEPC1.1 than other species. CREs of the 3kb of the 5’-flank regions of C_4_ genes were predicted applying the online tool Plantpan3.0 (score>=0.99). The TF families from top to bottom are ordered in a decreasing rank of total number of CREs from the five *Flaveria* species. (f) Word cloud represents the enriched CREs associated with C_4_ genes in the C_4_ species Ftri based on ATAC-seq, including those within 3kb upstream of start codon, within 3kb downstream of the stop codon and within the gene body.

We further characterized the distribution of CREs on the promoter regions of C_4_ genes. Ftri (C_4_) showed significantly more CREs for C_4_ genes than the other species did (p<0.001, *t*-test) (Figure 3d). Notably, when the total numbers of CREs of each gene in the five species were compared, ERF CREs were the most abundant of all examined C_4_ genes (Figure 3e and Supplemental Note 12). For example, for PEPC1 (PEPC1.1 from Ftri), there were 89 predicted ERF CREs from 5 species, followed by bZIP (54 CREs) and GATA (51 CREs).

To examine whether the ERF CREs were localized in the accessible chromatin regions (ACRs) in the C_4_ species, we analyzed the enriched CREs in the ACRs (ACR-CREs) obtained from two biological ATAC-seq (Supplemental Note 11). We categorized genes associated with ACRs-CREs into three types according to their distance to the nearest gene, *i.e.*, genic (gACR-CREs; overlapping a gene), upstream (upACR-CREs; within 3 kb upstream of the start codon of a gene) or downstream (downACRs-CRES; within 3 kb downstream of the stop codon of a gene). We then calculated enriched CREs in ACR-CREs. Across all three types of ACR-CREs, ERF CREs had the highest abundance among enriched CREs. Moreover, ERF dominated the enriched ACR-CREs of C_4_ genes, as well as in photosynthetic and photorespiratory genes (Figure 3f and Figure S4).

Taken together, our results suggested possible roles of both tandem duplications and recruitment of ERF CREs in the elevation of transcript abundances of C_4_ genes in the C_4_ species Ftri.

### Intronless ERF transcriptional factors were recruited in parallel in different C_4_ species

Given that many ERF CREs were recruited by C_4_ genes in Ftri (C_4_), we tested whether cognate ERF TFs were recruited in the same manner by C_4_ photosynthesis. We constructed a genome-wide co-regulatory network (GRN) of the five *Flaveria* species based on the gene expression profiles of at least 18 RNA-seq datasets either from a previous work (Zhu, 2020) or generated in the current study (Supplemental Note 13). We then obtained the sub GRN comprising C_4_ genes and their co-regulated TFs (C_4_GRN). TFs that had no predicted cognate CREs within 3 kb upstream of the start codon were filtered out. ERF, bHLH, MYB, NAC, and C2H2 were the top five most abundant TF families in the C_4_GRN of the five *Flaveria* species (Figure 4a and Figure S5). In the C_4_ species Ftri, 324 TFs were predicted to be co-regulated with C_4_ genes (Figure 4b), among which bHLH was the most prevalent TFs, with 29 genes, followed by the MYB related and ERF TF families, with 27 and 26 genes respectively (Figure 4c). Notably, ERF TFs were much more abundant in the C_4_GRN of the C_4_ species than in other species, though the number of predicted ERF TFs were comparable in all five *Flaveria* species (Figure 4a and Supplemental Note 13), suggesting that ERF TFs were preferentially recruited by C_4_ genes during evolution.

**Figure 4.**
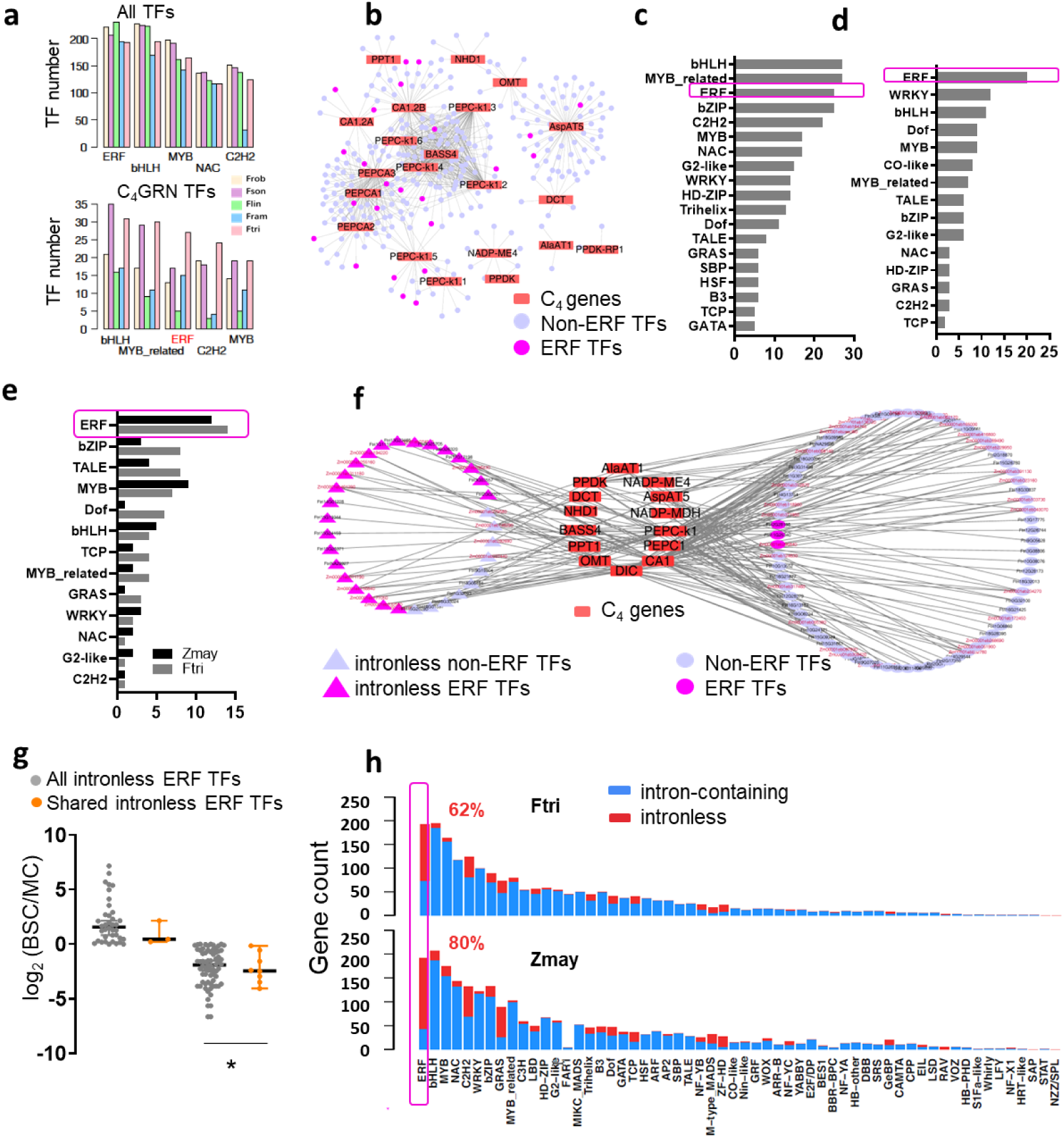
Intronless ERF were recruited by C_4_ genes in different C_4_ lineages. (a) The Top five most abundant TF families of all annotated TFs (top panel) and TFs that are co-regulated with C_4_ genes (bottom panel) (a) The network of C_4_ genes and TFs, which is termed as C_4_GRN in Ftri. (b) Families of TFs from C_4_GRN of Ftri. (c) Families of TFs from C_4_GRN of Zmay. Zmay C_4_GRN is extracted from published gene regulatory network in Tu et. al, 2019. (d) Orthologous TFs from C_4_GRN of Ftri and Zmay and their distribution in TF families. Orthologous groups were predicted with Orthofinder, and orthologous TFs between Ftri and Zmay, termed as shared TFs, are those from the same orthologous groups. (e) Regulatory network of shared TFs and C_4_ genes in Ftri and Zmay. Note that among the shared ERF TFs, 12 of 14 in Ftri and 11 of 12 in Zmay are intronless genes. (f) Bundle sheath cell (BSC) and mesophyll cell (MC) preferential expression of all intronless ERF TFs and shared intronless ERF TFs in Zmay. Y-axis shows log2 ratio of transcript abundance of each gene in BSC to that in MC. (g) Number of intronless genes in each TF family. (Abbreviations: GRN: gene co-regulatory network, MC: mesophyll cell, BSC: bundle sheath cell, Ftri: *Flaveria trinervia*, Zmay: *Zea mays*.)

C_4_ photosynthesis has appeared in more than 65 evolutionary independent lineages (Sage et al., 2012), and ERF CREs were previously found abundant in other C_4_ lineages, including *Zea mays* (corn; herein Zmay), *Setaria italica* (foxtail millet), and *Sorghum bicolor* (sorghum) (Supplemental Note 12) (Burgess et al., 2019; Marand et al., 2021). We investigated whether ERF TFs were also convergently recruited in other C_4_ lineages. We used Zmay, a model species for C_4_ research, for this test. Specifically, we analyzed a recently published leaf GRN for this species, which was constructed based on a combination of Chip-seq data, gene co-expression data, and a machine-learning based co-localization model(Tu et al., 2020). This genomic scale GRN included 1,475 TFs from 54 TF families, in which bHLH was the most prevalent family, with 138 genes, followed by ERF and MYB, with 136 and 108 genes, respectively (Tu et al., 2020) (Figure S6a). The C_4_GRN included 108 TFs from 15 TF families (Figure S6b), in which ERF TFs was the most prevalent ones, with 20 genes, followed by the WRKY and bHLH families, with 12 and 11 genes, respectively (Figure 4d). Therefore, ERF TFs were convergently recruited in both Ftri and Zmay, whose last common ancestor diverged around 160 mya (Kumar et al., 2017).

We further identified the shared TFs between Ftri and Zmay, *i.e.*, those recruited by both species and found in the same orthologous group based on Orthofinder’s analysis (Methods). Shared TFs were not required to regulate the same C_4_ genes in the two species. Our analysis found shared TFs from 27 orthologous groups which included 63 TFs from Ftri and 47 TFs from Zmay respectively. Again, the ERF TFs were the most abundant families in both species, including 14 (22.2%) and 12 (25.5%) of shared TFs in Ftri and Zmay, respectively (Figure 4e), and the targeted genes of these shared TFs covered 14 of the 15 C_4_ genes (Figure 4f). Notably, among the shared ERF TFs, 12 out of the 14 in Ftri and 11 out of 12 in Zmay were intronless genes, which account for 66.7% and 73.3% of total shared intronless TFs in Ftri (18 intronless TFs) and Zmay (15 intronless TFs), respectively (Figure 4f). Compared to other intronless ERF TFs in Zmay, the shared intronless ERFs showed more MC preferential expression (Figure 4g).

We further investigated the portion of intronless genes in each of the TF families in Ftri and Zmay. Intronless genes showed the most occurrences in ERF families, in which, 62% and 80% of ERF TFs were intronless in Ftri and Zmay respectively, accounting for 35.2% and 26.3% of the total intronless TFs in these species (Figure 4h). ERF TFs were also the most abundant intronless TF family in other land plant species, regardless of whether they are monocotyledonous or dicotyledonous, C_3_ or C_4_ species (Figure S7). These intronless ERFs showed greater changes in transcript abundances in response to low CO_2_ stress compared to intron-containing ERFs in the C_4_ species Ftri but not in other non-C_4_ *Flaveria* species. Similarly, intronless ERF TFs exhibited rapid and increased changes in gene expression in response to light induction in the C_4_ species Zmay but not in the C_3_ species *Oryza sativa, i.e.*, rice (*P*<0.05, Wilcoxon.test, Figure S8). These properties of intronless ERF TFs, including MC preferential expression (Figure 4g) and greater responses to low CO_2_ and light, might have contributed to their role in C_4_ photosynthesis.

We further analyzed the properties of one shared intronless ERF TF, *i.e.*, EREB34 (Figure S9a) between the Ftri C_4_GRN and Zmay C_4_GRN. EREB34 showed conserved expression profiles along the leaf developmental gradient or leaf age between C_3_ and C_4_ species. However, EREB34 showed significantly higher transcript abundance in the C_4_ species than in C_3_ species, which was shown both when we compared the evolutionarily close C_3_ and C_4_ species pairs individually (Figure S9b), and when we compared 30 C_4_ and 17 C_3_ species representing 18 independent lineages of C_4_ evolution (Steven Kelly, 2018) (Figure S9c). EREB34 also showed MC preferential expression in C_4_ species (Figure S9d). All these results suggested that EREB34 may play a role in the evolution of C_4_ photosynthesis.

### Intronless ERF TFs recruited by C_4_ photosynthesis originated in the Late Ordovician around 450 million years ago

ERF TFs belong to the AP2/ERF superfamily, which is one of the largest families of plant-specific TFs, and play vital roles in responses to various biotic and abiotic stresses (Feng et al., 2020; Gu et al., 2017; Xie et al., 2019). Our data showed that intronless ERF TFs were recruited as major regulators of C_4_ photosynthesis in both monocots and dicots. Considering that monocots and dicots diverged ∼160 mya (Kumar et al., 2017), while C_4_ photosynthesis emerged ∼ 35 mya (Sage et al., 2011), elements shared between the monocotyledonous and dicotyledonous C_4_ species were likely recruited before the divergence of monocots and dicots. We hence examined the origin of the intronless ERF TFs in plants. Specifically, we first surveyed the distribution of intronless genes in all annotated TF families based on the plantTFDB online tool (Jin et al., 2017) in 23 species spanning a wide spectrum of Viridiplantae (green plants), including four species from Chlorophyta, *Marchantia polymorpha* (Mploy, liverwort), which is regarded as one of the earliest land species (Delaux et al., 2019), seven monocotyledonous species, and 11 dicotyledonous species including the five *Flaveria* species sequenced here (Figure 5a). We included five and two C_4_ species from monocots and dicots, respectively.

**Figure 5.**
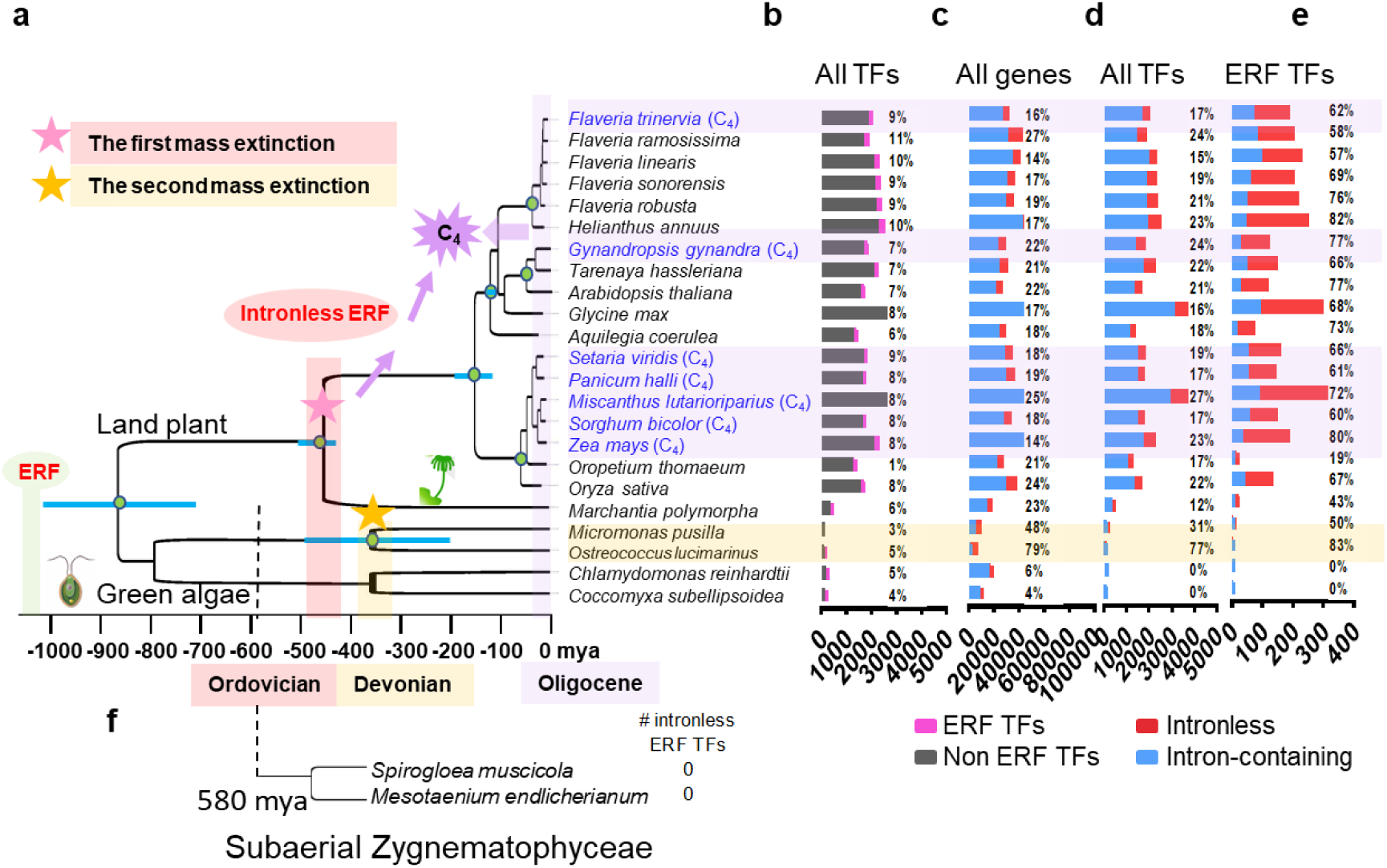
Intronless ERF TFs that were recruited in C_4_ photosynthesis emerged much earlier than C_4_ photosynthesis did. (a) Phylogenetic relationships of 23 species. C_4_ species are labeled in blue. The divergence time of each node is referenced from Timetree (http://timetree.org/). Two independent evolutionary origins of intronless ERF TFs are proposed, the occurrences of which coincide with the first mass extinction during the Late Ordovician (∼482 mya, pink bar) and the second mass extinction during the Late Devonian (∼358 mya, yellow bar), respectively. A star represents an independent evolutionary origin of intronless ERF TFs (b) Proportions of ERF TFs to total TFs. (c) Proportions of intronless gene to total protein coding genes. (d) Proportions of intronless TFs to total TFs. (e) Proportions of intronless ERF TFs to total ERF TFs. (f) The number of intronless ERF TFs in the two subaerial species from Zygnematophyceae, which is the sister branch of land plants and split from land plants ∼580 mya. (Abbreviation: ERF: ethylene responsive factor, mya: million years ago.)

We found that ERF TFs were present in all Viridiplantae, accounting for 7% of total TFs on average (Figure 5b). Furthermore, intronless genes were also present in all Viridiplantae, accounting for 22% of total annotated genes on average (Figure 5c). Intronless ERF TFs were found in all land plants and in the clade of Chlorophyta that includes *Micromonas pusilla* (Mpus), termed Mpus clade hereafter, but not the clade that includes *Chlamydomonas reinhardtii* (Crei), termed Crei clade hereafter (Figure 5d and Figure 5e). To determine whether the intronless ERF were specifically absent from the two species in the Crei clade or from the whole clade, we examined the other two species from the Crei clade with genome sequences available in the Phytozome database (https://phytozome-next.jgi.doe.gov), *i.e.*, *Dunaliella salina* and *Volvox carteri*. We found that intronless ERF TFs were not present in those two species either (Supplemental Note 14), implying that intronless ERF were absent from the Crei clade.

We then asked whether intronless ERF TFs were lost in Crei clade specifically or if there were two independent gains of intronless ERF TFs in the plant kingdom. We studied two species from Zygnematophyceae, which is the closest extant sister branch of land plants and evolved 580 mya (Gitzendanner et al., 2018), *i.e.*, *Spirogloea muscicola* and *Mesotaenium endlicherianum* (Cheng et al., 2019). Intronless ERF TFs were also absent in these two species (Figure 5f), suggesting that intronless ERF TFs in land plants and aquatic algae (Mpus clade) emerged from two independent evolutionary events. As further evidence, intronless ERF TFs from the Mpus clade showed nearly no orthologs in land plant species (Supplemental Note 14). Therefore, there were two independent gains of intronless ERF TFs during the evolution of Viridiplantae, and those evolved from the common ancestor of land plant species were recruited in C_4_ photosynthesis.

What might have promoted the emergence of intronless ERF TFs recruited to support C_4_ photosynthesis during the evolution of plants? To study this, we examined the recorded extreme climatic events around the period of the two independent occurrences of intronless ERF TFs in Viridiplantae. The first occurrence of intronless ERF TFs is around 450 mya when the land plants diverged from aquatic algae (Sanderson et al., 2004). This period coincided with the time of the Earth’s first mass extinction, around 447∼444 mya during the Late Ordovician (Finnegan et al., 2012; Sheehan, 2001). The second appearance of intronless ERF TFs, observed in the Mpus clade, occurred around 380 mya, which coincided with the time of the second mass extinction, around 372 mya during the Late Devonian (Da Silva et al., 2020; De Vleeschouwer et al., 2017) (Figure 5). Dramatic climate changes such as low temperature and oxygen deprivation have been proposed to underlie these two mass extinctions ^17,19^. Therefore, intronless ERF TFs might be the products of ancestral plants coping with extreme climate events. Long after the first emergence of these intronless ERF TFs in land species, around 35 mya (Sage et al., 2011), some of those TFs, especially those with strong cell specific expression patterns (Figure 4g) were recruited in C_4_ photosynthesis.

## Discussion

Identifying key regulators of C_4_ photosynthesis is a major task required for C_4_ engineering (Cui, 2021; Hibberd and Covshoff, 2010; Schluter and Weber, 2020; Westhoff and Gowik, 2010). The high-quality genome sequences of five *Flaveria* species offered a rich resource to support evolutionary and regulatory study of C_4_ photosynthesis. With this resource, we showed that intronless ethylene responsive factors (ERF) transcription factors (TF), a class of TFs involving in stress responses in plants (Christin et al., 2008; Ehleringer et al., 1991; Sage et al., 2011; Sage et al., 2012), played a role during the evolution of C_4_ photosynthesis. These intronless ERF TFs, originating from ∼450 mya (Sanderson et al., 2004) (Figure 5), have been repetitively recruited by different C_4_ lineages during evolution. Therefore, our results provided a molecular mechanism underlying shared TFs and *cis-*regulatory elements (CREs) between monocotyledonous and dicotyledonous C_4_ species that diverged 160 mya (Kumar et al., 2017), though the first C_4_ plants emerged ∼35 mya (Sage et al., 2011). The parallel recruitment of intronless ERF TFs implied that they may be used as targets during the current efforts in C_4_ engineering.

Why intronless ERF TFs? Intronless genes, featuring short mRNA length and lower transcript abundances compared to intron-containing genes (Shabalina et al., 2010) (Supplemental Note 15), play roles in plant responses to drought and salt stress (Liu et al., 2021). Intronless genes, regardless of being TF or not, showed more changes than intron-containing genes to low CO_2_ stress in all five *Flaveria* species, and greater and faster changes to light induction in both C_3_ and C_4_ species at the transcriptional level (Supplemental Note 15). Ethylene, an ancient plant hormone (Ju et al., 2015), bridges plant developmental adaptation and a changing environment (Merchante et al., 2013). ERF TFs, the last step of the ethylene signaling pathway, regulate the response of plants to environmental changes (Xie et al., 2019). Recently, one intronless ERF TF was reported to simultaneously modulate photosynthesis and nitrogen utilization in rice (Wei et al., 2022). ERF TFs showed remarkable changes to low CO_2_ stress in all five *Flaveria* species (Supplemental Note 15). Being evolutionary old and functioning in responding to environmental changes may underlie the observation that around 70% of ERF TFs evolved to be intronless genes in land plant species (Figure S7). In addition, ERF TFs existed widely across the plant kingdom, with a large presence of cognate CREs in plant genomes (Supplemental Note 12). The abundance of intronless ERF TFs and cognate CREs provided molecular resources for the evolution of C_4_ photosynthesis in coping with environmental stressors, such as low CO_2_, drought, and high light and high temperature conditions (Christin et al., 2008; Ehleringer et al., 1991; Sage et al., 2011; Sage et al., 2012) .

Intronless genes are also present in animals and fungi (see database: http://v2.sinex.cl/) (Jorquera et al., 2021). The ancient origin of intronless genes has been reported from animals. For example, intronless type I interferon (INF) in animals evolved from intron-containing type I INF in fish and amphibians around 350 mya during the Devonian (Gan et al., 2017), coinciding with the time when intronless ERF TFs originated in the Mpus algae clade. This suggested that environmental perturbations during the Devonian triggered the birth of new classes of intronless genes in both animals and plants. Interestingly, in humans, the counterpart of plant ERF TFs is the G-protein-coupled receptors (GPCRs). Around 50% of GPCRs are intronless genes, accounting for 53% of total human intronless genes (Gentles and Karlin, 1999; Grzybowska, 2012), compared to 70% ERF TFs that are intronless genes, accounting for around 30% of total plant intronless genes (Figure 5). Notably, ERFs in plants and GPCRs in humans have analogous functions in receiving and transducing signals from the external environment (Grzybowska, 2012; Xie et al., 2019). Therefore, particular types of intronless genes were retained for evolutionary adaptions in both the plant and animal kingdoms (Grzybowska, 2012; Xie et al., 2019).

## Acknowledgements

We appreciate Prof. Rowan F. Sage and Prof. Peter Westhoff for sharing us *Flaveria* materilas. The work is funded by Strategic Priority Research Program of the Chinese Academy of Sciences (grant number: XDB27020105), the general program of National Science Foundation of China (31870214) and the National Key Research and Development Program of China (2020YFA0907603).

## Authors’ contributions

XGZ, TL, CL and MJAL designed the study and wrote the paper. HD and ZG performed genome assembly and annotation, MJAL performed genome comparison analysis, qRT-PCR and RNA-seq analysis, HY conducted proteomics analysis, GC wrote the paper, FC performed gene regulatory network construction, YYZ performed PCR verification of the three paralogs of PEPC1s in Ftri, QT performed Ka/Ks analysis, FM and YW performed plasmodesmata analysis, CX performed transcriptional factor prediction, YZ and HL performed genome annotation of Fram, YT constructed *Flaveria* workspace in China National GeneBank (CNGB), LF and QG performed genome assembly of Fram, YQ perform transposon analysis, QZ and JZ performed syntenic analysis.

## Methods

### Plant materials and fluorescence in situ hybridization assay

*F. robusta* (Frob, C_3_) and *F. ramosissima* (Fram, C_3_-C_4_) were provided by Prof. Peter Westhoff (Heinrich Heine University, Germany). Seeds of *F. sonorensis* (Fson, C_3_-C_4_), *F. linearis* (Flin, C_3_-C_4_) and *F. trinervia* (Ftri, C_4_) were obtained from Prof. Rowan F. Sage (University of Toronto, Canada). Plants were grown in soil in green house as depicted in (Lyu et al., 2020).

The chromosome numbers of Frob, Flin and Ftri were investigated applying fluorescence in situ hybridization assay (FISH). Mitotic metaphase spreads of meristem root tip cells were prepared following (Deng et al., 2012). FISH was performed following (Li et al., 2019) with slight modifications, which is briefly depicted in Supplemental Note 2.

### Genome sequencing

Total DNA was extracted from young leaves. PacBio sequencing libraries were constructed following the tips of Pacific Biosciences (USA). DNA fragments of 0.5-18kb were chosen using BluePippin electrophoresis (Sage Science, USA). Libraries were then sequenced on the PacBio Sequel platform (PacBio, USA). The N50 of PacBio reads were from 16.4 to 21.9 kbp. Around 120 GB data were produced for each species on average. Genome coverage is from 66.9-fold (Ftri) to 232.2-fold (Frob). Besides, short reads were sequenced in Illumima X Ten platform in paired-end 150 bp mode. Around 200 million short reads were obtained for each species, which were used for genome assembly polishing as well as genome assembly completeness estimation. Hi-C libraries were constructed following (Mascher et al., 2017). Two Hi-C libraries were constructed for each species, with an inserted size of ∼350 bp, libraries were sequenced in Illumima X Ten platform. From 291Gb to 325Gb 150-bp paired-ended cleans data were generated for each species.

### De novo assembly

*Flaveria* nuclear genome sequences were assembled into 18 pseudochromosomes in a step-wise way. Sequencing adaptors were removed, and reads with low quality and short length were filtered applying PacBio SMRT Analysis package with following parameters: readScore, 0.75; minSubReadLength 50. The remained high-quality PacBio subreads were then corrected and contigs were assembled using Canu (v1.8) (Koren et al., 2017) with following parameters: useGrid = true, minThreads=4, genomeSize=1200m, minOverlapLength = 500, minReadLength = 1000. For contig polishing, the Illumina paired-end reads were mapped to assembled contigs applying bwa mem (bwa v0.7.17) (Li and Durbin, 2009), low qualified mapped reads were filtered off applying samtools (v1.11) (Li et al., 2009) with q30 setting. Pilon (v1.22) (Walker et al., 2014) were applied to polish with the following parameters: --mindepth 10 --changes --fix bases.

For Fram specifically, the BioNano next-generation mapping system was used to help high-quality genome assembly. DNA was labelled at Nt.BspQI sites applying the IrysPrep kit (BioNano Genomics, USA). Molecules collected from BioNano chips (BioNano Genomics, USA) were de novo assembled applying RefAligne and Assembler offered on the BioNano (Pendleton et al., 2015) using following parameters: -U -d -T 20 -j 4 -N 10 -i 5, which resulted in the optical genome maps. Next, genome assembly resulting from Pilon (v1.22) (Walker et al., 2014) mentioned above were then evaluated and corrected by aligning with the optical genome maps. Corrected contigs and optical genome maps were aligned and merged applying hybridScaffold.pl (Pendleton et al., 2015) which resulted in hybrid scaffolds. Next, HERA(Du and Liang, 2019) was used to fill gaps of obtained hybrid scaffold in following parameters: InterIncluded_Side=30000, InterIncluded_Identity=99, InterIncluded_Coverage=99, MinIdentity=97, MinCoverage=90, MinLength=5000, MinIdentity_Overlap=97, MinOverlap_Overlap=1000, MaxOverhang_Overlap=100, MinExtend_Overlap=500. Obtained hybrid scaffolds were then used for following assembly.

Followed, assembled genome sequences were improved using Hi-C data in two steps. First, contigs were corrected using Hi-C data. Briefly, low-quality Hi-C data (over 10% N base pairs or Q10 < 50%) were removed, and remained reads were mapped to assembled contigs applying bwa (v0.7.17) (Li and Durbin, 2009) with ‘aln’ settings and other parameters were in default. Only uniquely mapped reads were used to perform re-assembly. Invalid mapping was filtered off applying HiC-Pro (v2.11.1) (Servant et al., 2015) with following settings: mapped_2hic_fragments.py -v -S -s 100 -l 1000 -a -f -r -o. Next, corrected contigs were re-assembled into scaffold applying LACHESIS(Burton et al., 2013) with following parameters: CLUSTER MIN RE SITES = 770, CLUSTER MAX LINK DENSITY=2, CLUSTER NONINFORMATIVE RATIO = 2, ORDER MIN N RES IN TRUNK=578, ORDER MIN N RES IN SHREDS=593.

### Annotation of transposable elements

To predict transposable elements (TEs), whole genome sequences of the five *Flaveria* species were searched for repetitive sequences individually. A de novo repeat sequence library was constructed by RepeatModeler (RepeatModeler-Open-1.0.5) with the following parameters: RepeatModeler -database database_name -engine ncbi -pa [int]. Then, we used RepeatMasker (RepeatMasker-Open-4.1.0) to search for similar TEs against the de novo library with the following parameters: RepeatMasker genome.fa -lib de_novo_library -nolow -no_is -q -engine rmblast -pa [int] –norna. Intact long terminal repeat retrotransposons (LTR-RTs) were identified using LTR_FINDER (v1.07) (Xu and Wang, 2007) and LTRharvest (v1.5.10) (Ellinghaus et al., 2008) with the default parameters. And Then LTR_Retriever (v2.9.0) (Ou and Jiang, 2018) was used to merge the above results with the parameters: LTR_retriever -genome genome.fa -inharvest species.harvest.scn -infinder species.finder.scn –nonTGCA species.harvest.nonTGCA.scn. The insertion time of intact LTR-RT was extracted from LTR-Retriever analysis.

### Annotation of protein coding genes

Gene models were predicted by combining de novo prediction, homology-based and transcriptome-based strategies. Briefly, Augustus (v2.4) (Stanke and Morgenstern, 2005), GlimmerHMM (v3.0.4) (Majoros et al., 2004), GeneID (v1.4) (Parra et al., 2000) and Genscan (http://genes.mit.edu/GENSCAN.html) were used in combination for de novo prediction. GeMoMa (v1.3.1) ^(Keilwagen et al., 2019)^ was used for homology-based prediction. To facilitate gene annotation, from 18 to 32 Illumina RNA-seq datasets were generated either in this study (for Flin, as depicted below) or generated in our previous work (Zhu, 2020). Clean RNA-seq reads were mapped to genome applying Hisat2 (v2.0.4) (Kim et al., 2019) and genome-based transcript assembly was performed applying StringTie (v1.2.3) (Pertea et al., 2015) in default parameters. Besides, de novo transcript assembly was conducted based on RNA-seq data applying PASA (v2.0.2) (Haas et al., 2003) in default parameters. All predicted gene structures were integrated into consensus gene models using EVidenceModeler (v1.1.1) (Haas et al., 2008), and pseudo genes were predicted applying GeneWise (v2.4.1) ^(Birney et al., 2004)^, Coding sequence (CDS) failed to be translated either lacking an open reading frame (ORF) or having premature stop codons were removed.

The completeness of protein repertoire was estimated in different aspects: 1) using BUSCO (v3.0.2) (Seppey et al., 2019) against to viridiplantae reference, 2) RNA-seq reads mapping to genome applying STAR (v2.7.3a) (Dobin et al., 2013), and 3) 150-bp paired-ended DNA sequencing reads mapping to genome apply bowtie2 (v2.3.4.3) (Langmead and Salzberg, 2012) (Supplemental Note 3).

Putative gene functions were assigned using the best match to GO, KEGG, Swiss-Prot, TrEMBL and a non-redundant protein database (NR) using BLASTP (v2.2.31+) (Camacho et al., 2009) with the E value threshold of 1e-5.

Transcriptional Factors were predicted using online website PlantTFDB (v5.0) (Jin et al., 2017; Tian et al., 2020) (http://planttfdb.gao-lab.org/prediction.php). *Cis*-regulatory elements (CREs) of promoter regions (3kb upstream of the start codon) were predicted using Plantpan (v3.0) (Chow et al., 2019) with a score threshold of 0.99.

### Orthologous genes prediction and gene evolution

To predict orthologous groups, protein coding genes from the five *Flaveria* species, *Arabidopsis thaliana* (Atha), *Helianthus annuus* (Hann, sun flower), and *Lactuca sativa* (Lsat, lettuce) were predicted applying Orthofinder (v2.3.11) (Emms and Kelly, 2019) using default parameters. The protein sequences of Atha (TAIR10), Hann (v1.0) and Lsat (v7) were downloaded from Phytozome (v13) (https://phytozome.jgi.doe.gov/pz/portal.html). In case where there were multiple alternative transcripts, the longest one was kept to represent the protein-coding gene.

### Phylogeny and divergence time analysis

To construct the phylogenetic tree, CDS sequences of 1:1 orthologous genes were aligned applying MUSCLE (v3.8.31) (Edgar, 2004) in default parameters. Alignments of all the CDS were linked to make a super matrix, and RAxML (v7.9.3) (Stamatakis, 2006) was then applied for inferring phylogenetic tree using the following model: GTR (General Time Reversible nucleotide substitution model) + GAMMA (variations in sites follow GAMMA distribution) + I (a portion of Invariant sites in a sequence). To calibrated the evolutionary time, CDS were aligned codon-wisely guided by protein alignment using pal2nal (v14) (Suyama et al., 2006).The evolutionary time was calibrated applying mcmctree in PAML package (v4.9) (Yang, 2007) using the following parameters: seqtype=0 (nucleotides), clock=2 (independent), model = 0 (JC69). The reported fossil time between Hann and Lsat, *i.e.*, 34∼40 million years as inferred from timetree (http://timetree.org/) was used for calibration. The phylogenetic tree and calibrated evolutionary time were displayed using FigTree (http://tree.bio.ed.ac.uk/software/Figuretree/).

### Synteny analysis between *Flaveria* species

To identify syntenic gene blocks in each species and between Frob with other four species, all-against-all BLASTP (E value < 1e−10, top five matches) (v2.2.31+) (Camacho et al., 2009) was performed for protein coding genes for each genome pairs. Syntenic blocks were determined according to the presence of at least five synteny gene pairs applying MCScanX (v0.8) (Wang et al., 2012) with default parameters. Colinearity of the five species were drawn with JCVI (https://github.com/tanghaibao/jcvi). Circular graphic was plotted using Circos (v0.69-5).

### Investigation of light responsiveness of C_4_ genes using qRT-PCR

Consider that C_4_ genes showed fast light responsiveness in C_4_ species but not in C_3_ species (Burgess et al., 2016; Lyu et al., 2020), to verify the identified C_4_ version of C_4_ genes, we investigated the changes of gene expression in response to light induction using quantitative real time PCR (qRT-PCR). *Flaveria* species were put to dark room at 6:00 pm. The dark-adapted plants were illuminated at 9:00 am the next day. Fully expanded leaves, usually the 2^nd^ or 3^rd^ leaf pair counted from the top, were cut after the leaves were illuminated for different time periods, *i.e.*, 0, 2, and 4 h, and then flashed into liquid nitrogen. Samples were stored at −80°C before processing. RNA isolation and qRT-PCR were performed as described earlier in(Lyu et al., 2020). Relative transcript abundances were calculated by comparing to ACTIN7, the primers used here were as depicted in our previous work (Zhu, 2020) .

### RNA-seq and transcriptional quantification for *Flaveria* species

RNA-seq data of Flin were obtained from plant grown under low CO_2_ (100 ppm) *vs* normal CO_2_ (380 ppm) for two weeks and four weeks respectively, and plant grown under high light (with PPFD of 1400 μmol m^-2^ s^-1^) *vs* control light condition (500 μmol m^-2^ s^-1^) were sequenced independently. Growth conditions were as depicted in (Zhu, 2020). For RNA extraction, the young fully expanded leaf usually situated on the 2^nd^ or 3^rd^ pair of leaves counting started from the top was used. The chosen leaves were cut and immediately frozen into liquid nitrogen and stored thereafter at -80 °C until further processing. Total RNA was then isolated following the protocol of the PureLInk^TM^ RNA kit (ThermoFisher Scientific, USA). The RNA-sequencing was performed in the Illumina platform in the paired-end mode with a read length of 150 bp. RNA-seq data of other four species were from our previous work (Zhu, 2020).

To quantify the expression level of *Flaveria* genes, raw reads were trimmed applying fastp (v0.20.0) (Chen et al., 2018) using default parameters. Transcript abundance of gene were calculated by mapping RNA-seq reads to assembly genome sequence of corresponding species using RSEM (v1.3.3) (Li and Dewey, 2011) in default parameters, where STAR (v2.7.3a) (Dobin et al., 2013) was selected as the mapping tool.

### Proteomics

Mature leaves were cut from one-month old plant as depicted above, and leaves were put into liquid nitrogen quickly. Frozen leaf samples were grinded thoroughly and then incubated in lysis buffer (50 mM ammonium bicarbonate, 8M urea, 1mM DTT, complete EDTA-free protease inhibitor cocktail (PIC) (Roche)). Samples were centrifuged at 14,000g for 10 min at 4 °C. The supernatant was kept for total protein samples. Total protein concentration was measured with a Bradford assay (Bradford, 1976).

The process of protein digestion, HPLC Fractionation, LC-MS/MS analysis and data processing were detailed in Supplemental Note 8. Briefly, to generate data dependent acquisition (DDA) library, peptides were prefractionated. Fractionated peptides were mixed from all the 30 samples (a total of 200 μg). The mixture was separated by a linear gradient, and finally, 30 fractions were mixed into 15 components. Raw data from each species were used to construct library based on protein sequence from such species. As a result, five peptide libraries were obtained with one for each species. Finally, data independent acquisition (DIA) was performed using Spectronaut (version 14.7, Biognosys, Zurich, Switzerland). Default settings for quantification at MS1 level were employed for quantification. The mass spectrometry proteomics data have been deposited to the PRteomics IDEntifications Database (PRIDE).

### ATAC-seq for the C_4_ species Ftri

To isolate nuclei from C_4_ species Ftri, fully expanded mature leaves were harvest at 1:00 pm. Around 3g fresh leaves from 5 plants were used for each of the two biological replicates. Leaf materials were grinded in ice in 10 ml 4xNE buffer (40 mM MES -KOH, PH5.4, 40 mM NaCl, 40 mM KCl, 10mM EDTA, 1M Sucrose, 0.1 mM spermidine, 0.5mM spermine and 1mM DTT). Next, the debris was removed by sieving through two layers of 70 μm nylon cell strainer into precooled flasks and then the fluid were centrifuged at 200g at 4 °C for 3min to further remove debris. The supernatant was centrifuged at 2000g at 4 °C for 5min to spin down Nuclei. Nuclei were lysed by adding 1X NE buffer, 0.1% (v/v) NP40 and 0.1 (v/v) Tween-20 and incubated on ice for 3 min. Nuclei were pelleted by centrifugation at 2000g at 4°C for 5 min. Pellets were then incubated in RS buffer (Tn5 mix, 10 mM Tris-HCL, PH 7.4, 10 mM NaCl, 3mM MgCl_2_, 0.01% digitonin, 0.1% OM and 0.1% Tween-20) at 37 °C for 30 min. The Tn5 tagmentation was then terminated under 95 °C for 2 min. DNA was purified using a spin column (Qiagen, Germany) and then amplified using index primers matching the Illumina Nextra adapter.

ATAC-seq libraries containing DNA insert between 50 and 150 bp were gel purified and sequenced in Illumima X Ten platform in paired-end 150 bp mode. Raw reads were trimmed using fastp (v0.20.0) (Chen et al., 2018) in default parameters. Sequencing reads were mapped to genome sequence of Ftri (C_4_) using bowtie2 (v2.3.4.3) (Langmead and Salzberg, 2012) in default parameters. Mapping result were sorted using “sort” function in samtools (v1.11) (Li et al., 2009), and mapping with low quality was filtered off using “view” function in samtools with -q=10. Duplicated mapped reads were removed using “rmdup” function in samtools. Mapping peaks were then called using macs2 (v2.2.7.1) (Zhang et al., 2008) using the following parameters: -f BAMPE, -g 1.7e9 -q 0.05, --broad --nomodel --min-length 50. The parameter “broad” was used to allow closed peaks merging into a broad peak. We referred peaks predicted in this study as Tn5 hyper sensitive site (THS).

Peaks associated genes were assessed using “closest” function in bedtools (v2.29.2) (Quinlan and Hall, 2010) with -k 2, considering the closest two genes (both upstream and downstream). The distribution of THS relative to genome feature were assessed using “computeMatrix” function in deeptools (v3.5.0) (Ramirez et al., 2014) with the following parameters: --skipZeros –reference Point TSS -a 3000 -b 3000, the result was then plotted using “plotHeatmap” in the same tool. To predict known motif of the THS, the function fimo within meme package (v5.0.2) (Grant et al., 2011) was applied to scan known motifs annotated in Plantpan 3.0 (Chow et al., 2019) through the sequences of THS with default parameters.

### Comparison of intron-containing and intronless genes in *Flaveria* and other species

To calculate proportions of intronless genes in different species, we classified intronless genes in different species based on gene annotations. A gene was classified as intronless gene if all of its transcriptional isoforms contains exact one exon. For non-*Flaveria* species studied here, genome sequences, gene annotation files and protein sequences were downloaded from accessible databases. Briefly, those of Zea mays (v5) were downloaded from Maize GDB (https://maizegdb.org/), those of *Spirogloea muscicola* and *Mesotaenium endlicherianum* were downloaded from figshare (figshare.com) referencing from (Cheng et al., 2019), and those of *Miscanthus lutarioriparius* (Mlut) was downloaded from figshare referencing from (Miao et al., 2021), and those of the rest species were downloaded from Phytozome (v13) (https://phytozome-next.jgi.doe.gov/), with genome versions as following: Atha (TAIR10), *Hann* (v1.2), *Glycine max* (v2.0), *Aquilegia coerulea* (v3.1), *Oropetium thomaeum* (v1), *Setaris viridis* (Svir, v2.1), *Panicum Hallii* (Phal, v2.1), *Sorghum bicolor* (Sbic, v3.1.1), Osat (v7), *Marchantia* polymorpha (v3.1), *Panicum halli* (v2.0), *Chlamydomonas reinharditii* (Crei, v5.5), *Micromonas* pusilla (v3.0), *Coccomyxa* subellipsoidea (v2.0), *Dunaliella* salina (v1.0), *Ostreococcus* lucimarinus (v2.0) and *Volvox carteri* (v2.1).

To compare the transcript abundance of intron-containing and intronless genes for non-*Flaveria* species, and to compare the expressional pre ferences of intron-containing and intronless genes in mesophyll cells, we either inferred transcript abundances of genes from published references or calculated gene expression levels based on published RNA-seq datasets as detailed in Supplemental note 14. Specifically, we thank Eric Schranz (Wageningen University), Andreas Weber (Heinrich Heine University) and Julian Hibberd (Cambridge University) for access to the *Ggyn* genome sequence and the updated Thas genome assembly (Cheng et al., 2013) for classifying intronless genes and performing the RNA-seq quantification.

### Data availability

The genome assemblies, gene annotations, transcriptome data, proteomics data and raw reads are available at China National GeneBank (CNGB) (https://db.cngb.org/codeplot/datasets/public_dataset?id=flaveria) with project ID of CPN0003058. The genome assemblies, gene annotations, transcriptome data, proteomics data are also available at figshare (https://figshare.com/account/home#/projects/114567). The genome assemblies are also available at National Center for Biotechnology Information (NCBI) with accession number SAMN14943594 for *F. robusta*, SAMN14943595 for *F. sonorensis*, SAMN14943597 for *F. linearis*, SAMN14943596 for *F. ramosissima* and SAMN14943598 for *F. trinervia*. The mass spectrometry proteomics data were submitted to PRoteomics IDEntifications Database (PRIDE) with accession number PXD024720 (username, password: M6E7WzlM). RNA-seq data of Flin were submitted to Gene Expression Omnibus (GEO) in the NCBI database available with accession number: PRJNA827625. RNA-seq data of Frob, Fson, Fram and Ftri are from published data with project accession PRJNA600545.

## Supplemental information

1. **Supplemental Notes:** including supplemental note 1 ∼ supplemental note 20, which contain methods and results that support the main conclusion of the work.

2. **Supplemental Table and Figures**: including Table S1 and Figure S1 ∼ Figure S9.

## Supplemental Table and Figures

**Table S1.**
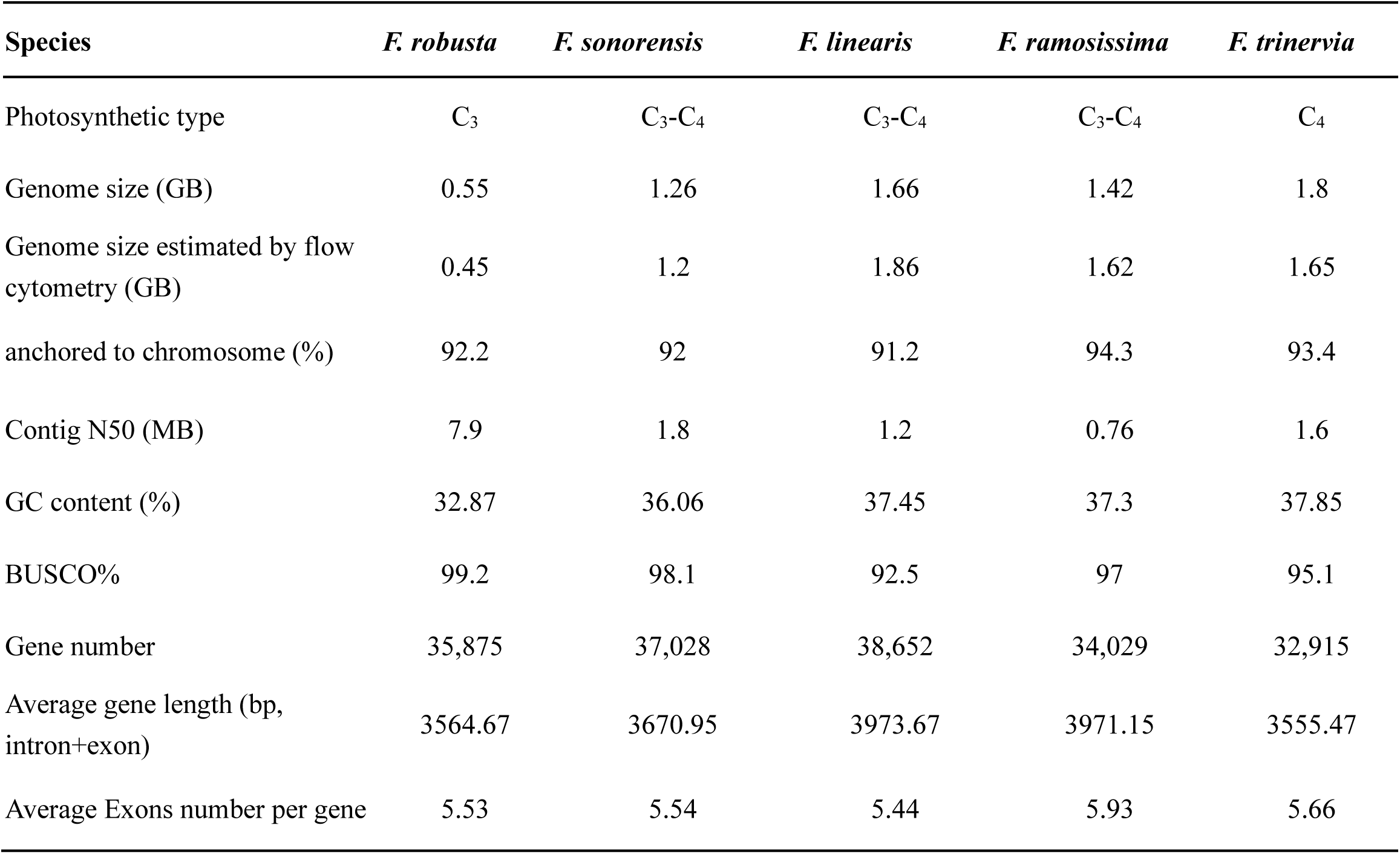
Statistics of genome assemblies and annotations.

**Figure S1.**
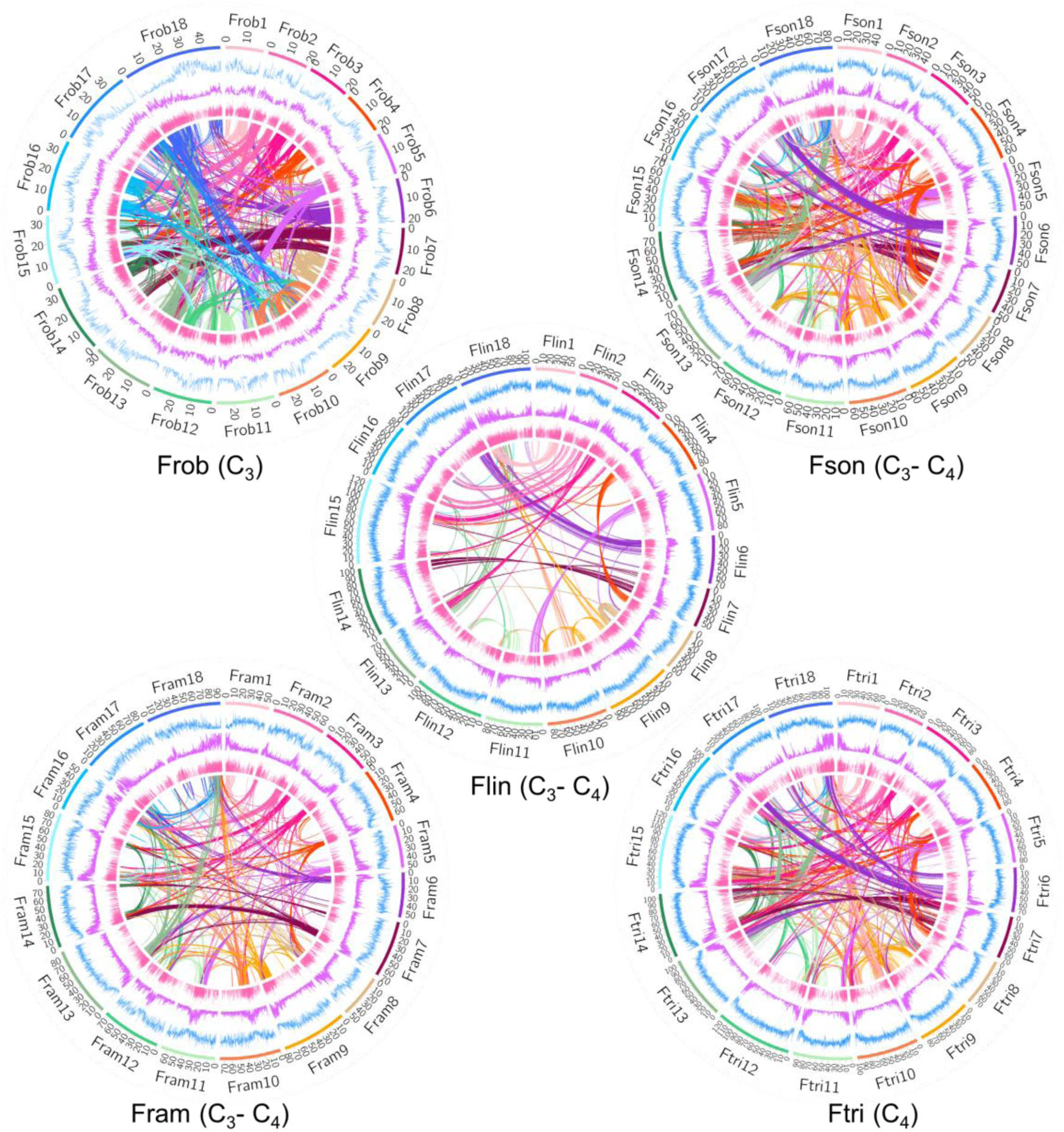
Genome features of five *Flaveria* species. The circular representation of pseudochromosomes. From outer to inner side: blue: LTR density per million base pair (Mb), purple: exon density per Mb, pink: transcript abundance per gene in log10 TPM (transcript per million mapped reads). Lines in the inner circle represent links between synteny-selected paralogs. (Abbreviations: Frob: *F. robusta*, Fson: *F. sonorensis*, Flin: *F. linearis*, Fram: *F. ramosissima*, Ftri: *F. trinervia*)

**Figure S2.**
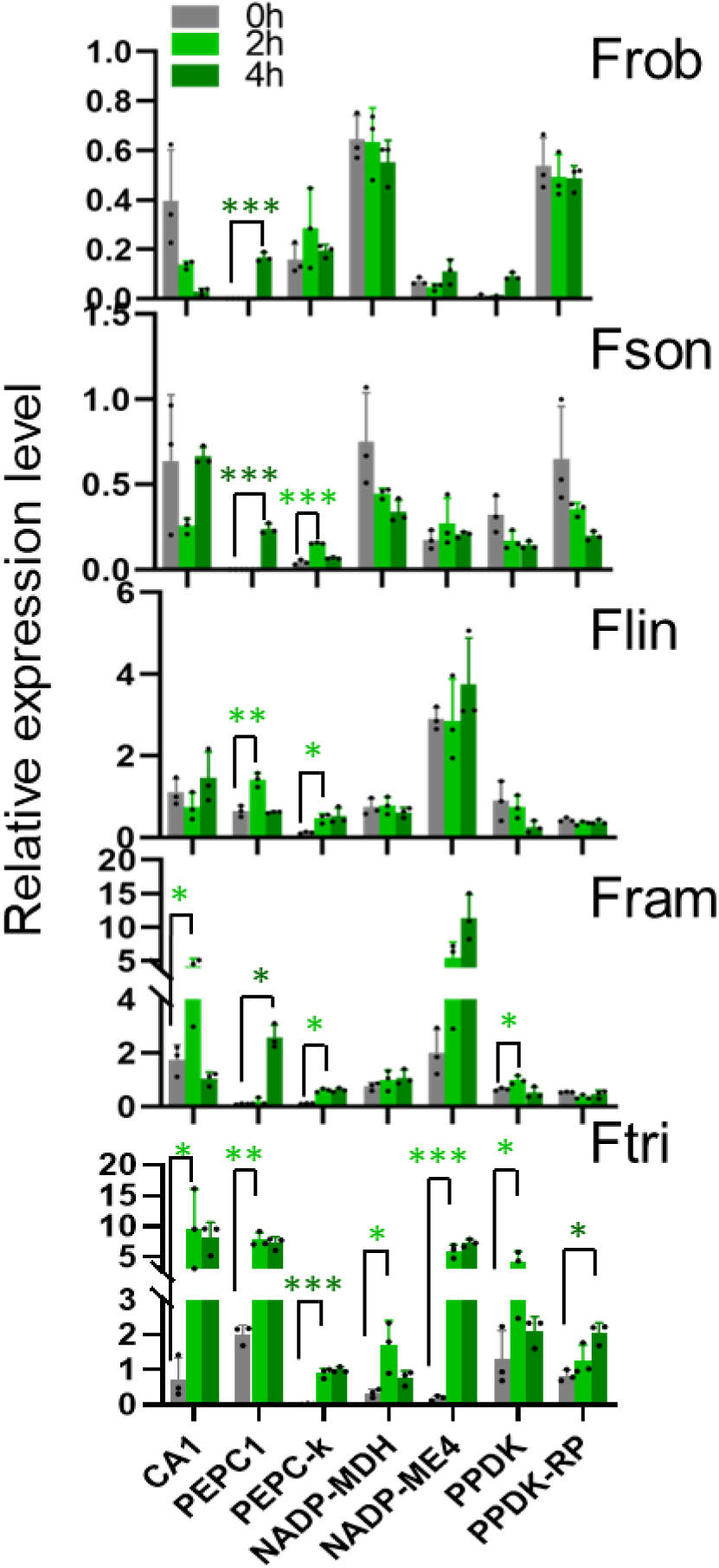
C_4_ gene gradually gained light responses during evolution. Real-time quantitative (qRT)-PCR was used to quantify the transcript abundance of C_4_ enzymes in mature leaves after 0, 2 and 4h upon illumination. significance levels were calculated using *t*-test. (*: 0.05–0.01, **: 0.01–0.001, ***: < 0.001) (Abbreviations: CA1, carbonic anhydrase 1; PEPC1, phospho*enol*pyruvate carboxylase 1; PEPC-k: PEPC kinase; NADP-MDH, NADP-dependent malate dehydrogenase; NADP-ME4, NADP-dependent malic enzyme 4; PPDK, pyruvate/orthophosphate dikinase; PPDK-RP, PPDK regulatory protein)

**Figure S3.**
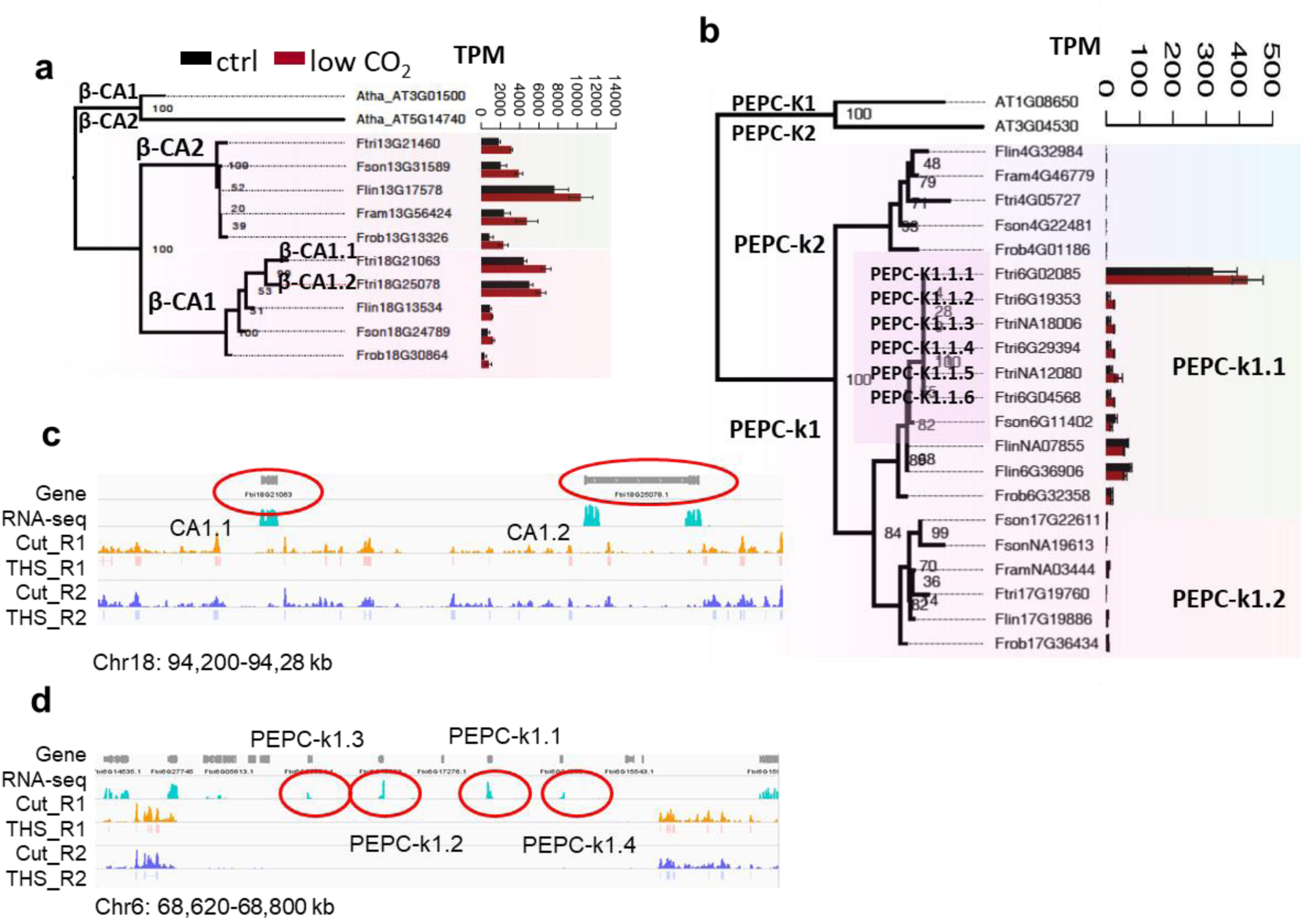
C_4_ version of CA and PEPC-k show more copies in the C_4_ species Ftri resulting from tandem duplication. (a) and (b) illustrate the gene tree of CA and PEPC-k respectively. Gene tree were constructed based alignment of protein sequences. Bootstrap scores were from 100 bootstrap samplings. Bars show gene expressions of leaves from two-month-old plants grown in low CO_2_ condition (100 ppm) *vs* normal CO_2_ condition (380 ppm) for four weeks. Three biological replicates were performed for each condition. (c) and (d) Integrated Genome Viewer (IGV) of RNA-seq reads and ATAC-seq reads of two copies of CA1 and four copies of PEPC-k1 anchored to chromosomes in Ftri respectively. Tn5 cuts and transposase hypersensitive sites (THS) from two biological replicates are showed. (Abbreviations: CA1: carbonic anhydrase1; PEPC-k1: phospho*enol*pyruvate carboxylase kinase1.)

**Figure S4.**
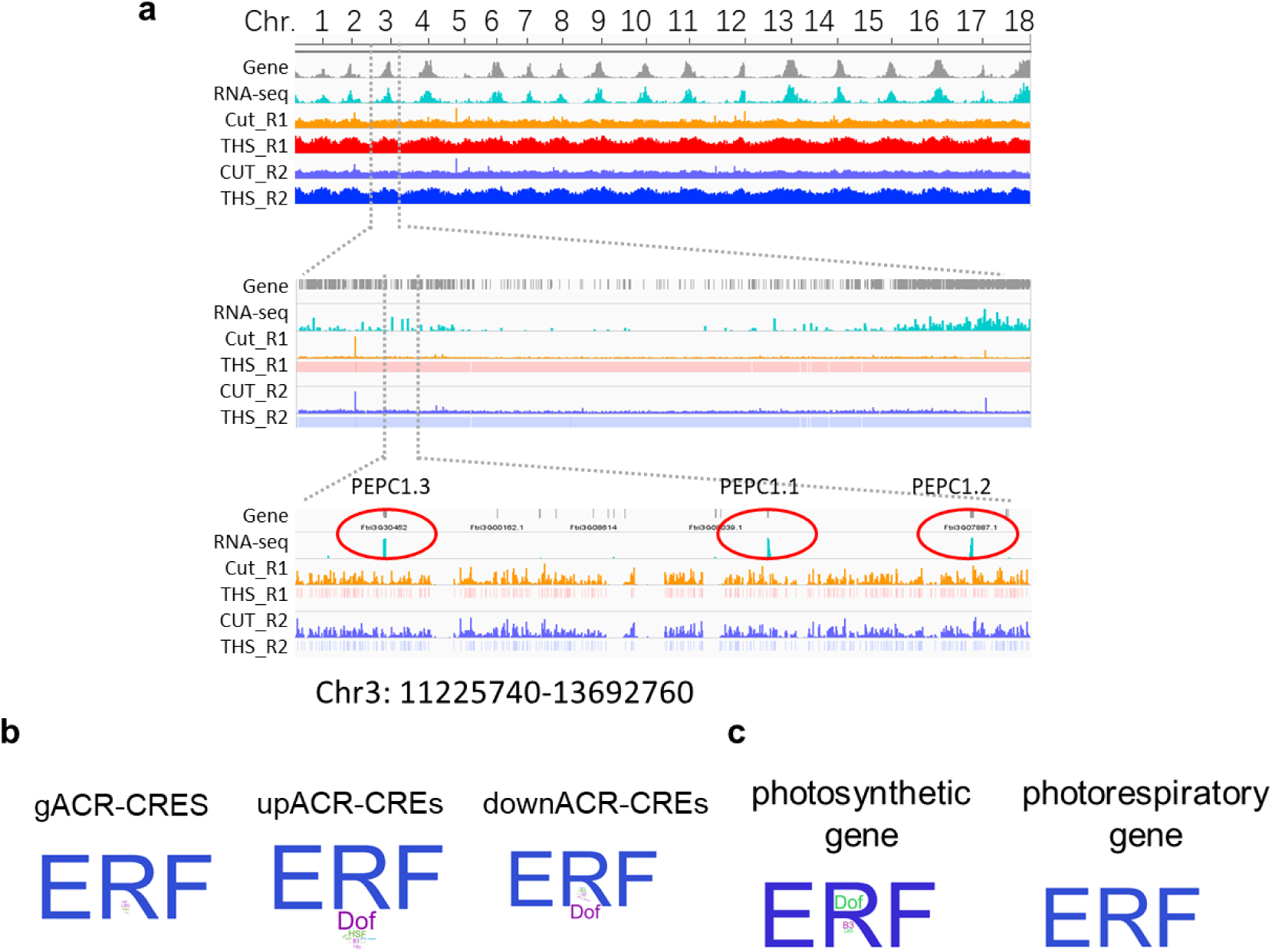
Predicted *cis*-regulatory elements in the C_4_ species Ftri by applying ATAC-seq. (a) Integrated Genome Viewer (IGV) of RNA-seq reads and two-biological replicates of ATAC-seq reads in Ftri are showed in three spatial resolutions, *i.e.*, genome scale with chromosome number and location indicated (top), chromosomal scale (middle), and million-base genomic region including PEPC1. (b) Enriched *cis*-regulatory elements (CREs) in three types of accessible chromatin regions (ACR-CREs), *i.e.*, genic (gACR-CREs: overlapping a gene), upstream (upACR-CREs: within 3kb upstream of the start codon of a gene) and downstream (down ACRs-CRES: within 3kb downstream of the stop codon of a gene). (c) Enriched ACR-CREs associated with photosynthetic genes and photorespiratory genes. (Abbreviations: ATAC-seq: transposase-accessible chromatin using sequencing; ACR: accessible chromatin regions; CREs: *cis*-regulatory elements)

**Figure S5.**
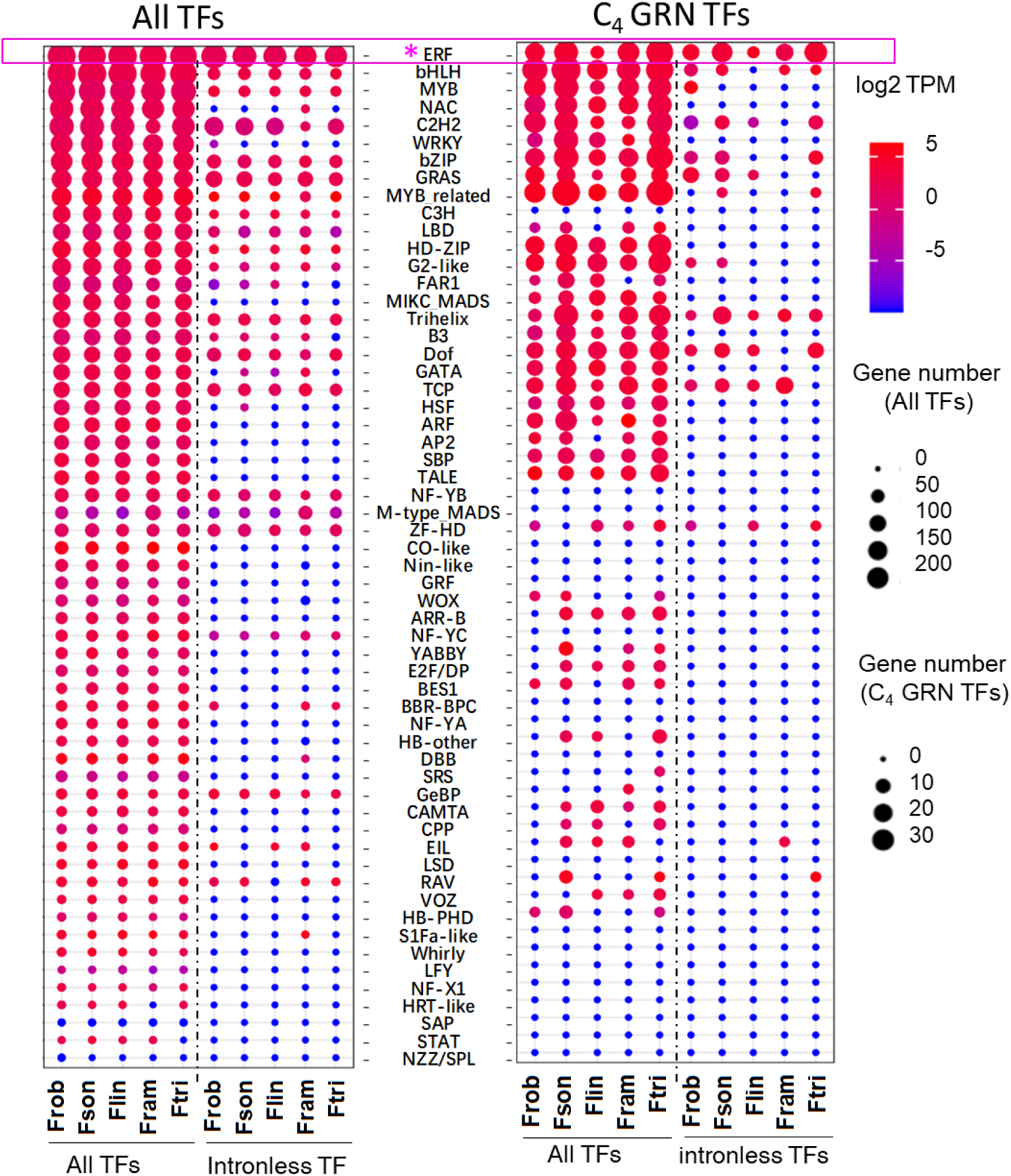
The number and transcript abundances of intron-containing and intronles genes in each TF family. Heat maps show the gene number and transcript abundance of genes in each TF family from all the annotated TFs (left panel) and from C_4_GRN (right panel). The size of circle represents the number of genes, and the color represents the log2 transformed transcript abundances in transcript per million mapped reads (TPM).

**Figure S6.**
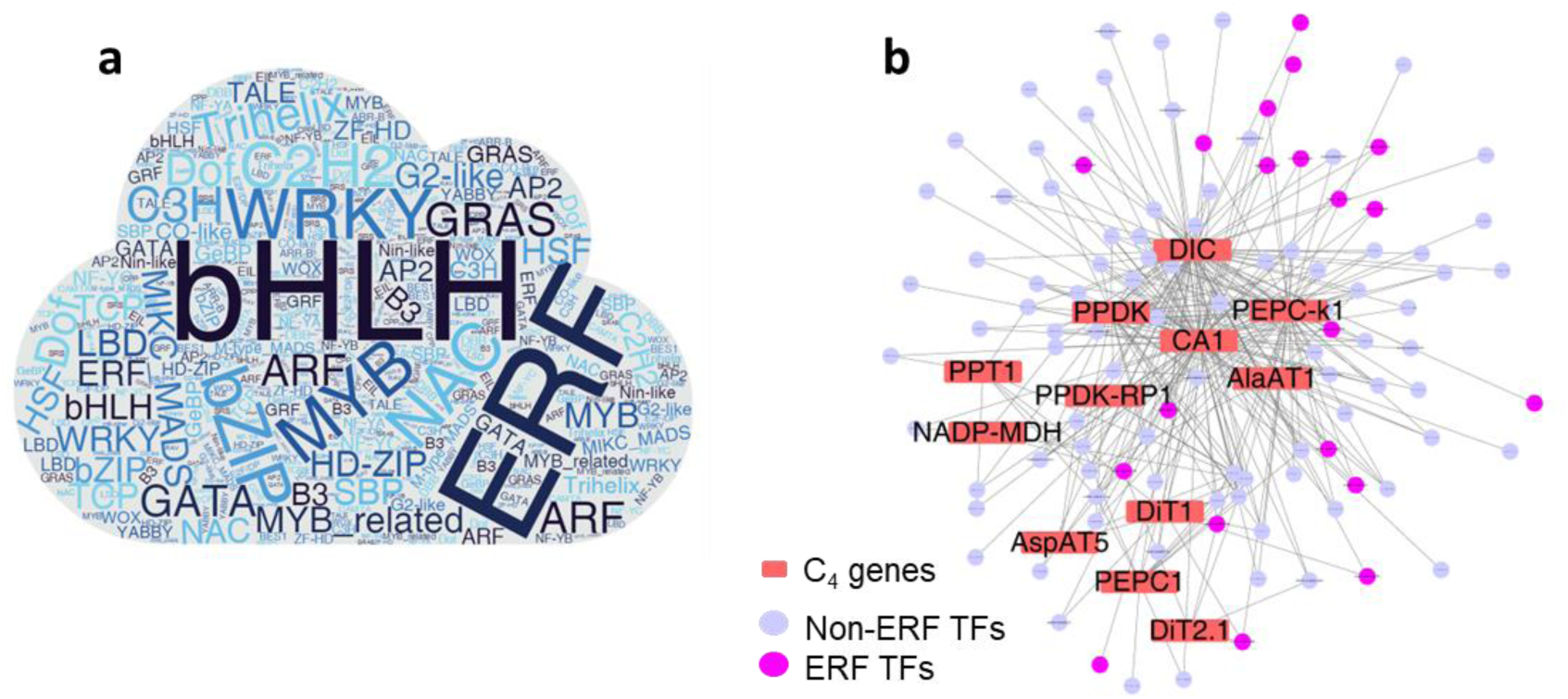
ERF TF were recruited by C_4_ genes in *Zea mays*. (a) Word cloud shows the frequencies of TFs in each TF families in the leaf gene regulatory network (GRN) of *Zea mays* (Zmay, corn) from (Tu et al., 2020). In the whole leaf GRN, bHLH is the most prevalent TFs with 138 genes. (b) The C_4_GRN in Zmay that includes C_4_ genes and their regulatory TFs. ERF is the most abundant TFs in the C_4_GRN as showed in purple circle. 12 C_4_ gene are included within the whole leaf GRN. (Abbreviation: Zmay: *Zea mays*)

**Figure S7.**
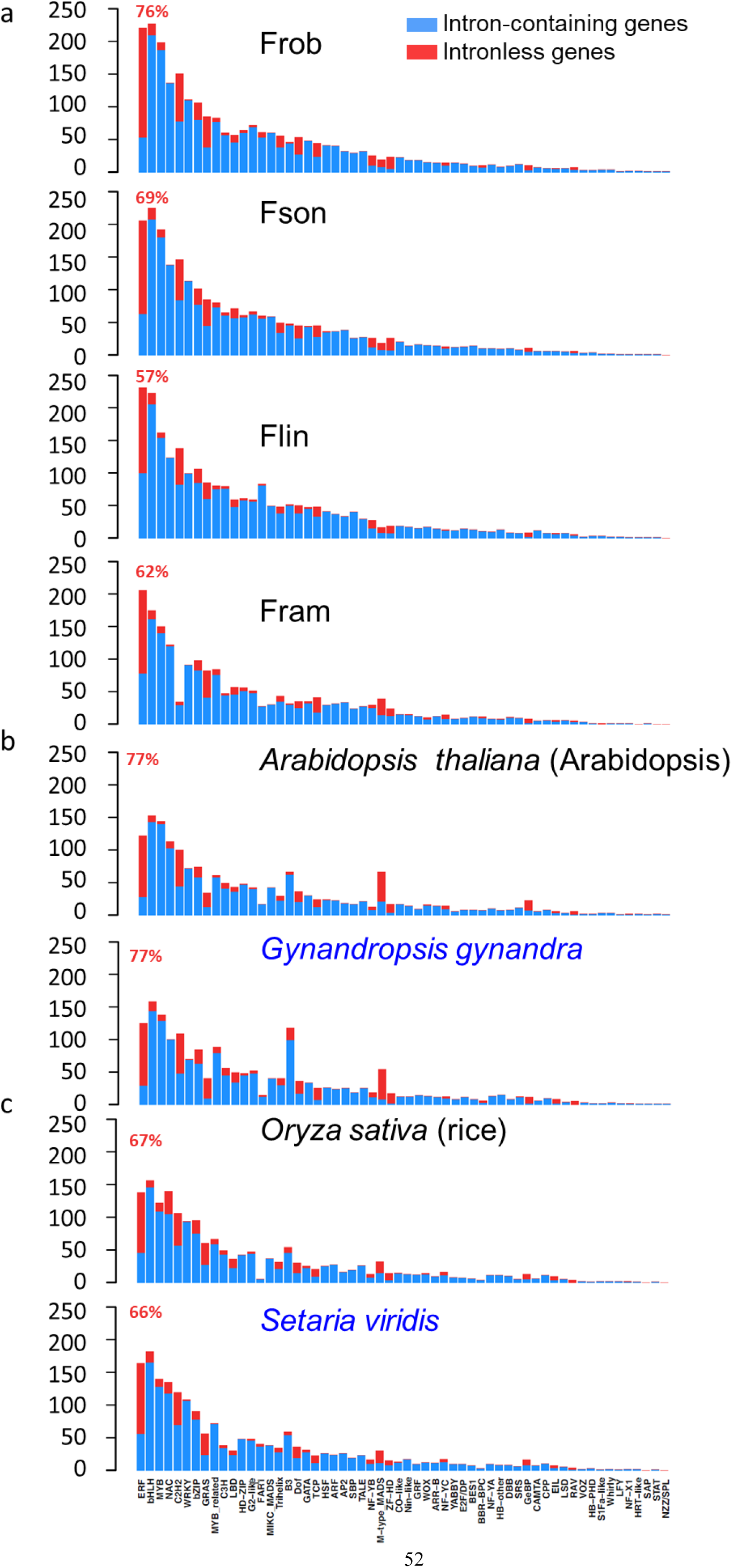
The number of intronless genes in each TF family in different species. The number of intronless gene and intron-contain genes are showed in each TF family from (a) four *Flaveria* species, (b) two dicotyledonous species and (c) two monocotyledonous species. C_4_ species are labeled in blue font. Proportions of intronless genes in ERF TF family are showed in red fond for each species.

**Figure S8.**
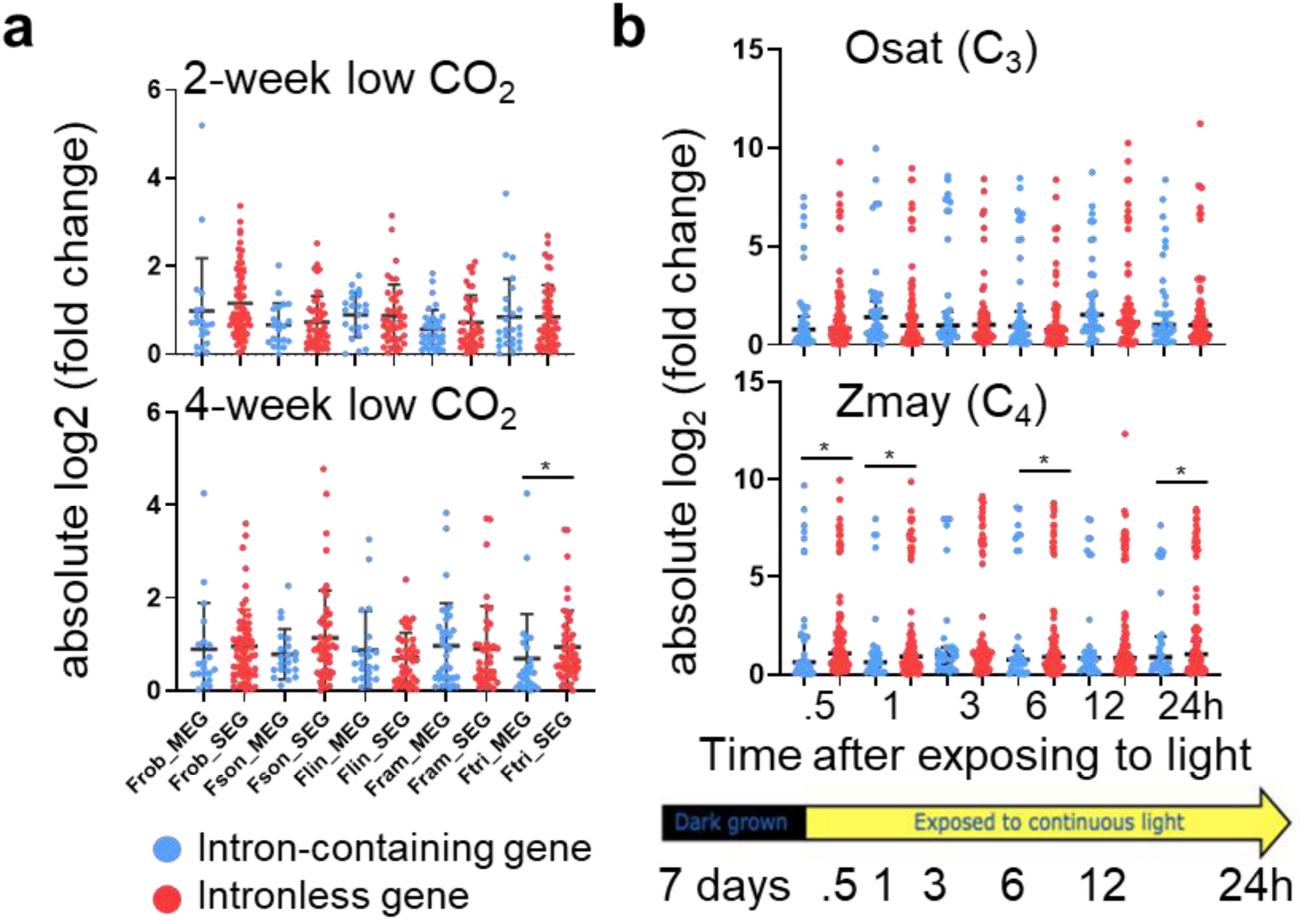
The response of intronless and intron-containing ERF TFs to environmental changes. (a) The changes on transcript abundances of intronless and intron-containing ERF TFs in five *Flaveria* species in response to low CO_2_ (100 ppm) compared to normal CO_2_ (380 ppm). RNA-seq data of both low CO_2_ and normal CO_2_ grown plants were taken from leaves after plants being grown under each condition for two weeks and four weeks respectively. (b) The response of transcript abundances of intronless and intron-containing ERF TFs in *Oryza sativa* (Osat, C_3_) and *Zea mays* (Zmay, C_4_) under light induction. The gene expression data of Osat and Zmay are from (Xu et al., 2016). Seeds of both species were germinated and grown under dark for 7 days. RNA-seq data of leaves were taken before light, 0.5h, 1, 3h, 6h, 12h and 24h after light respectively. The fold change of each time point was calculated as the ratio of gene expression level of this time point to that of the prior time point. Gene expression levels for all analysis was showed in transcript per million mapped reads (TPM). (Abbreviations: MEG: multi-exon genes, *i.e.*, intron-containing genes; SEG: single exon genes, *i.e.*, intronless genes; Osat: *Oryza sativa*; Zmay: *Zea mays*.)

**Figure S9.**
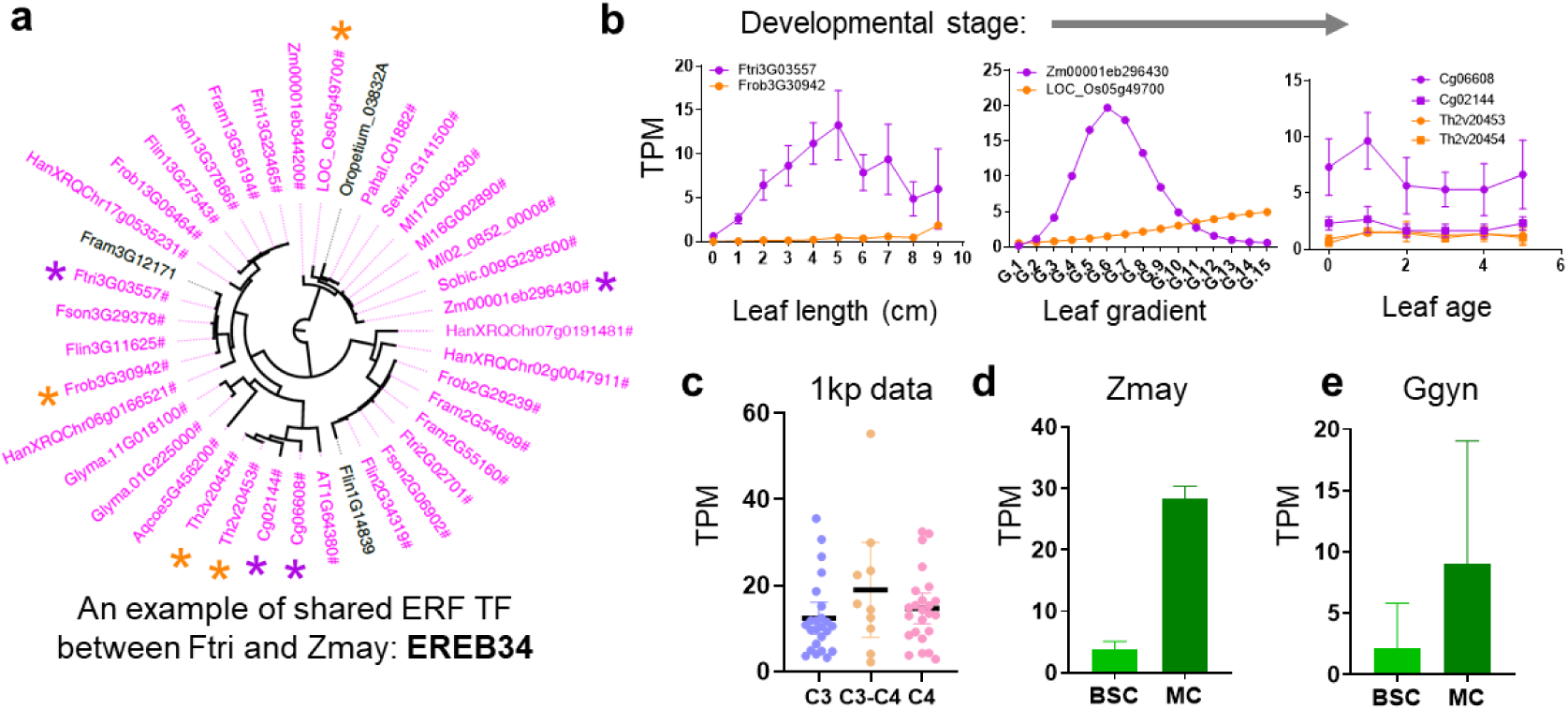
An example of intronless ERF TF that was recruited to regulate C_4_ genes in both Ftri and Zmay. (a) Gene tree of EREB34. Intronless genes are labeled in purple. EREB34 orthologous genes among 23 species (see Figure5) were predicted applying Orthofiner. EREB34 present in higher plants but not in algae or liverwort. Genes marked with orange and purple stars are from C_3_ and C_4_ plants that compared in transcript abundances in **(b)**. **(b)** Comparisons of EREB34 in transcript abundances between Frob (C_3_) *vs* Ftri (C_4_) (lef), *Oryza sative* (Osat, C_3_) *vs Zea mays* (Zmay, C_4_) (middle), and *Tarenaya hassleriana* (Thas, C_3_) vs *Gynandropsis gynandra* (Ggyn, C_4_) (right) along leaf developmental gradient. Genes from C_3_ species are labeled in orange, and those from C_4_ species are in purple. RNA-seq data of these species are from published sources, *i.e.*, data of Frob and Ftri are from (Billakurthi. et al., 2020), data of Osat and Zmay are from (Xu et al., 2016), data of Thas and Ggyn are from (Kulahoglu et al., 2014) . The first points in Frob and Ftri (0) represent meristem. Leaf ages of Thas and Ggyn are as following: **0**: 0-2 days (d); **1**: 2-4 d; **2**: 4-6 d; **3**: 6-8 d; **4**: 8-10 d and **5**: 10-12 d. **(c)** Transcript abundances of EREB34 in C_3_, C_3_-C_4_ and C_4_ species from one thousand plants (1kp) project, covering 18 independent C_4_ lineages. RNA-seq data are from (Steven Kelly, 2018). **(d)** Transcript abundance of EREB34 in Zmay bundle sheath cell (BSC) and mesophyll cell (MC). Expressional data are from (Chang et al., 2012). **(e)** Transcript abundances of EREB34 in Ggyn BSC and MC. Expressional data are from (Aubry et al., 2014). (Abbreviations: MC: mesophyll cell; BSC: bundle sheath cell.)

