## Supplementary Methods and Figures for "Born with intronless ERF transcriptional factors: C_4_ photosynthesis inherits a legacy dating back 450 million years"

### Supplemental Table and Figures

**Table S1. Statistics of genome assemblies and annotations**

| Species | *F. robusta* | *F. sonorensis* | *F. linearis* | *F. ramosissima* | *F. trinervia* |
|---|---|---|---|---|---|
| Photosynthetic type | $C_3$ | $C_3$-$C_4$ | $C_3$-$C_4$ | $C_3$-$C_4$ | $C_4$ |
| Genome size (GB) | 0.55 | 1.26 | 1.66 | 1.42 | 1.8 |
| Genome size estimated by flow cytometry (GB) | 0.45 | 1.2 | 1.86 | 1.62 | 1.65 |
| anchored to chromosome (%) | 92.2 | 92 | 91.2 | 94.3 | 93.4 |
| Contig N50 (MB) | 7.9 | 1.8 | 1.2 | 0.76 | 1.6 |
| GC content (%) | 32.87 | 36.06 | 37.45 | 37.3 | 37.85 |
| BUSCO% | 99.2 | 98.1 | 92.5 | 97 | 95.1 |
| Gene number | 35,875 | 37,028 | 38,652 | 34,029 | 32,915 |
| Average gene length (bp, intron+exon) | 3564.67 | 3670.95 | 3973.67 | 3971.15 | 3555.47 |
| Average Exons number per gene | 5.53 | 5.54 | 5.44 | 5.93 | 5.66 |

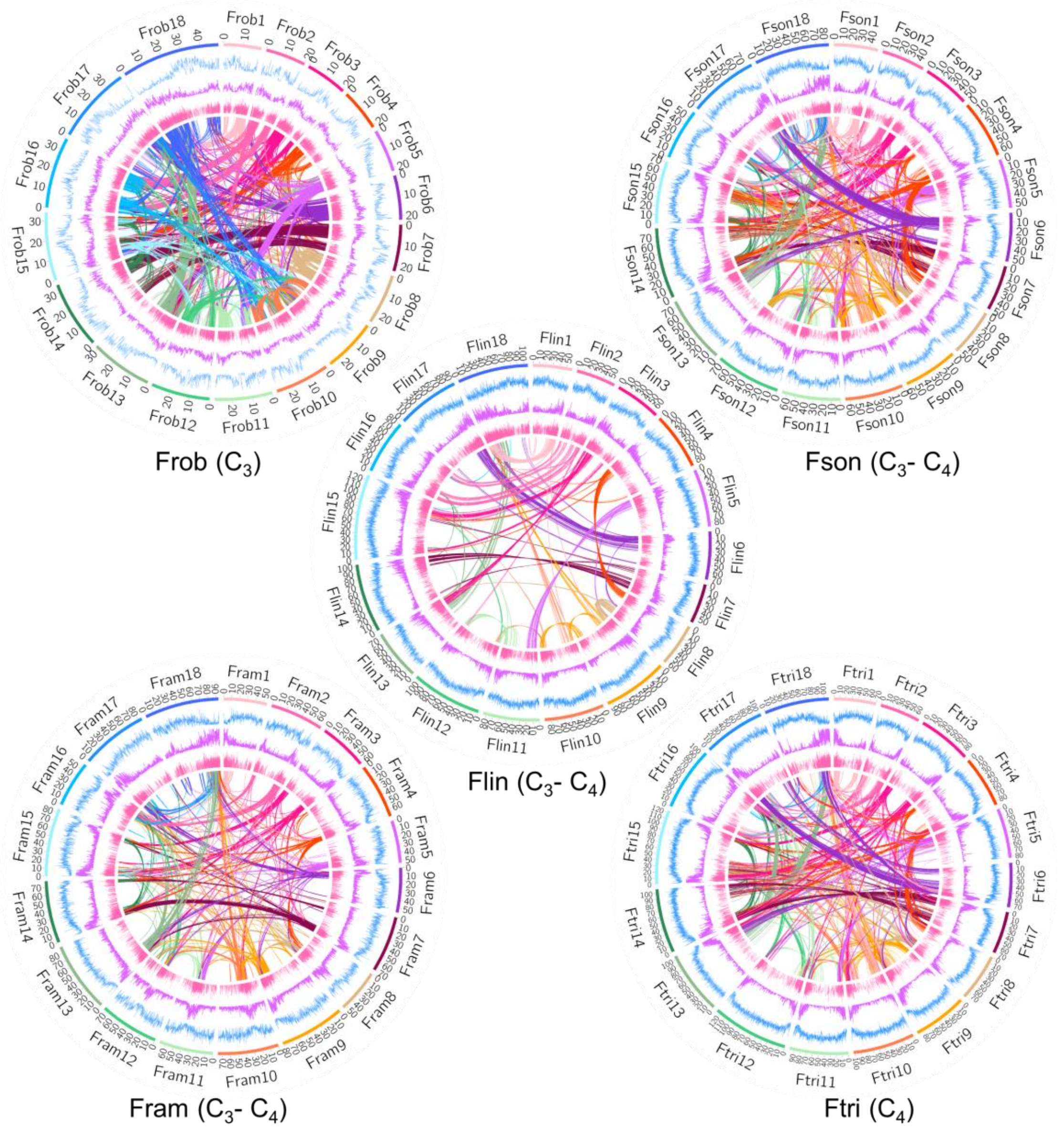


**Figure S1. Genome features of five *Flaveria* species**

The circular representation of pseudochromosomes. From outer to inner side: blue: LTR density per million base pair (Mb), purple: exon density per Mb, pink: transcript abundance per gene in log10 TPM (transcript per million mapped reads). Lines in the inner circle represent links between synteny-selected paralogs. (Abbreviations: Frob: *F. robusta*, Fson: *F. sonorensis*, Flin: *F. linearis*, Fram: *F. ramosissima*, Ftri: *F. trinervia*)

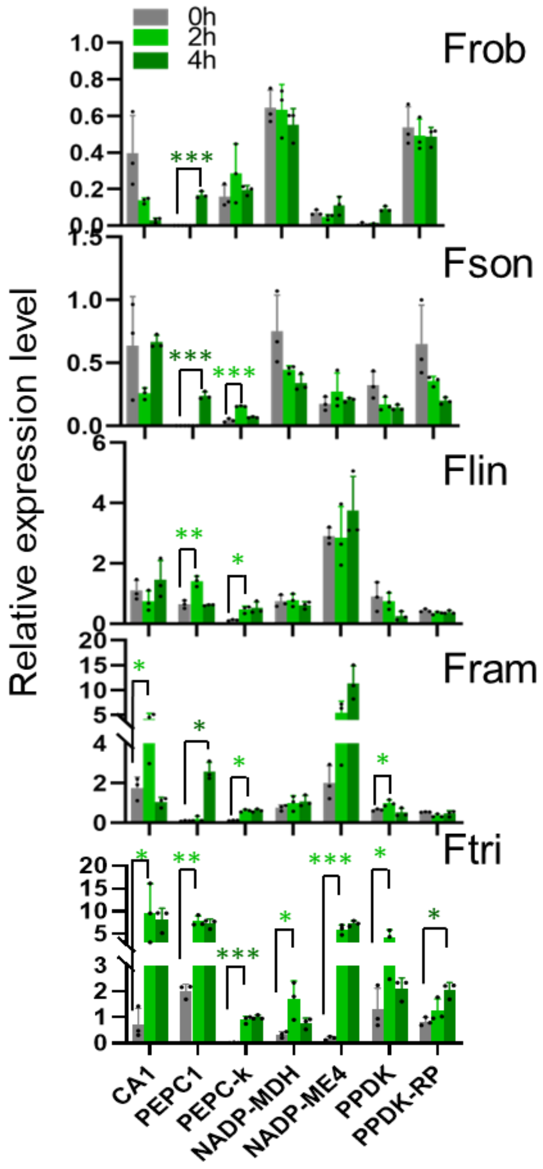


**Figure S2. $C_4$ gene gradually gained light responses during evolution**

Real-time quantitative (qRT)-PCR was used to quantify the transcript abundance of $C_4$ enzymes in mature leaves after 0, 2 and 4h upon illumination. significance levels were calculated using *t*-test. (*: 0.05–0.01, **: 0.01–0.001, ***: < 0.001) (Abbreviations: CA1, carbonic anhydrase 1; PEPC1, phospho*enol*pyruvate carboxylase 1; PEPC-k: PEPC kinase; NADP-MDH, NADP-dependent malate dehydrogenase; NADP-ME4, NADP-dependent malic enzyme 4; PPDK, pyruvate/orthophosphate dikinase; PPDK-RP, PPDK regulatory protein)

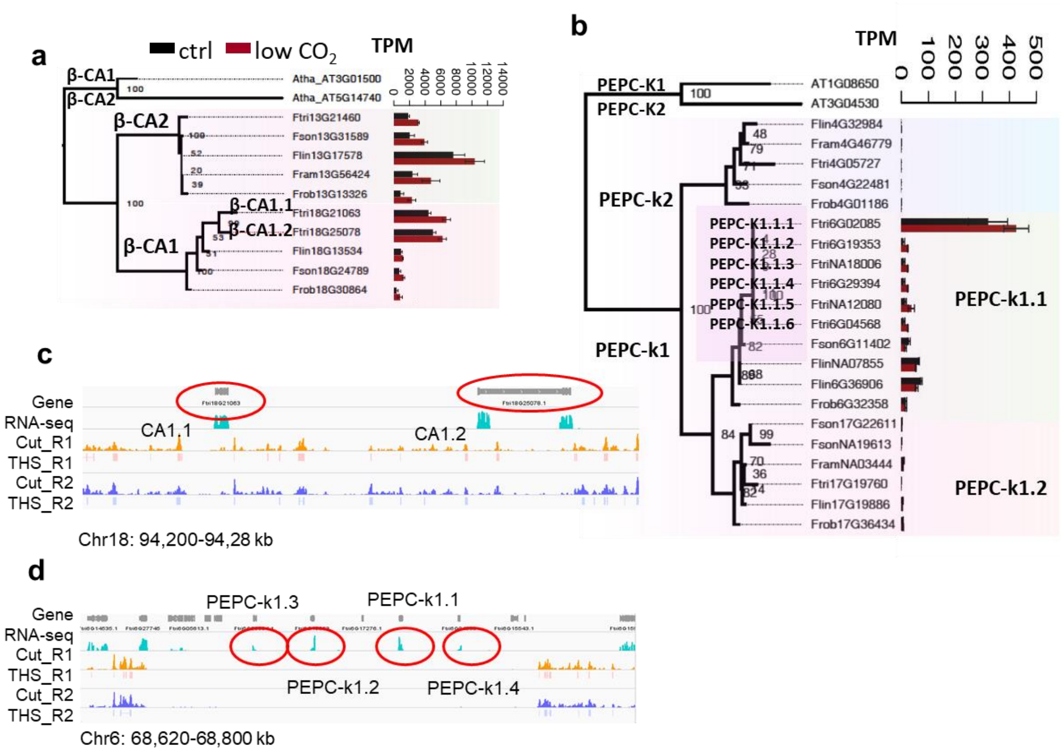


**Figure S3. C4 version of CA and PEPC-k show more copies in the C4 species Ftri resulting from tandem duplication**

(a) and (b) illustrate the gene tree of CA and PEPC-k respectively. Gene tree were constructed based alignment of protein sequences. Bootstrap scores were from 100 bootstrap samplings. Bars show gene expressions of leaves from two-month-old plants grown in low $CO_2$ condition (100 ppm) *vs* normal $CO_2$ condition (380 ppm) for four weeks. Three biological replicates were performed for each condition. (c) and (d) Integrated Genome Viewer (IGV) of RNA-seq reads and ATAC-seq reads of two copies of CA1 and four copies of PEPC-k1 anchored to chromosomes in Ftri respectively. Tn5 cuts and transposase hypersensitive sites (THS) from two biological replicates are showed. (Abbreviations: CA1: carbonic anhydrase1; PEPC-k1: phospho*enol*pyruvate carboxylase kinase1.)

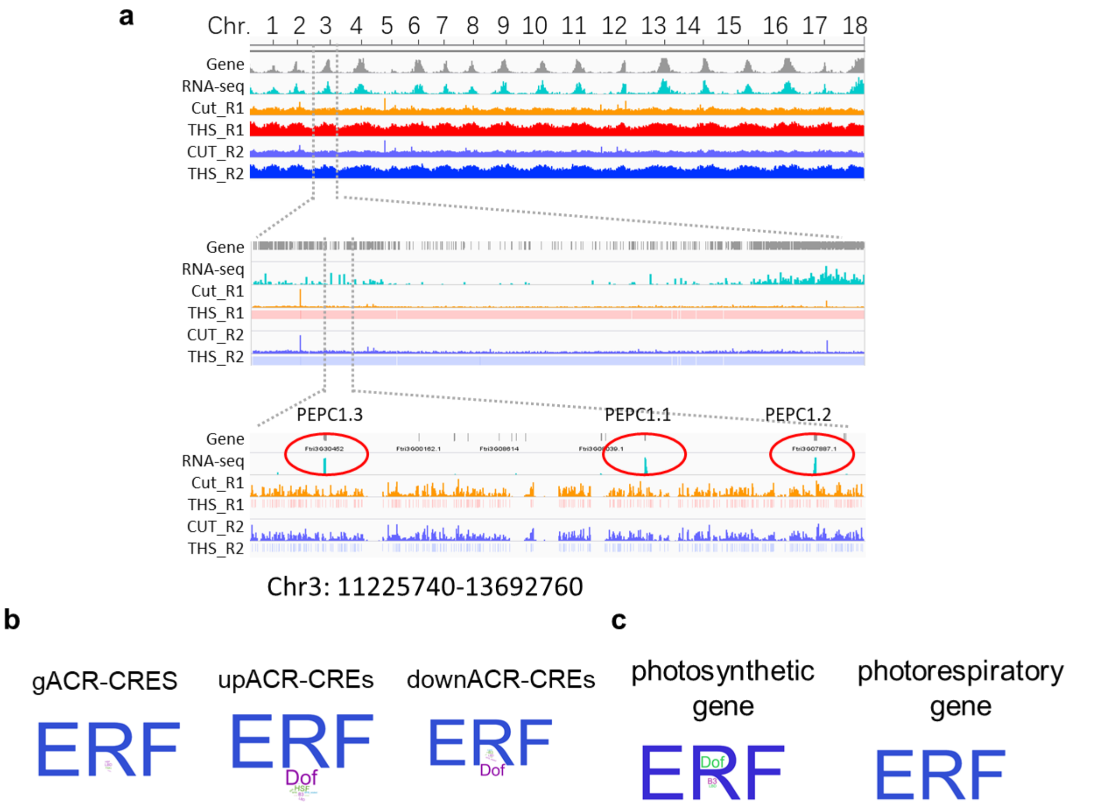


**Figure S4. Predicted *cis*-regulatory elements in the $C_4$ species Ftri by applying ATAC-seq**

(a) Integrated Genome Viewer (IGV) of RNA-seq reads and two-biological replicates of ATAC-seq reads in Ftri are showed in three spatial resolutions, *i.e.*, genome scale with chromosome number and location indicated (top), chromosomal scale (middle), and million-base genomic region including PEPC1. (b) Enriched *cis*-regulatory elements (CREs) in three types of accessible chromatin regions (ACR-CREs), *i.e.*, genic (gACR-CREs: overlapping a gene), upstream (upACR-CREs: within 3kb upstream of the start codon of a gene) and downstream (down ACRs-CRES: within 3kb downstream of the stop codon of a gene). (c) Enriched ACR-CREs associated with photosynthetic genes and photorespiratory genes. (Abbreviations: ATAC-seq: transposase-accessible chromatin using sequencing; ACR: accessible chromatin regions; CREs: *cis*-regulatory elements)

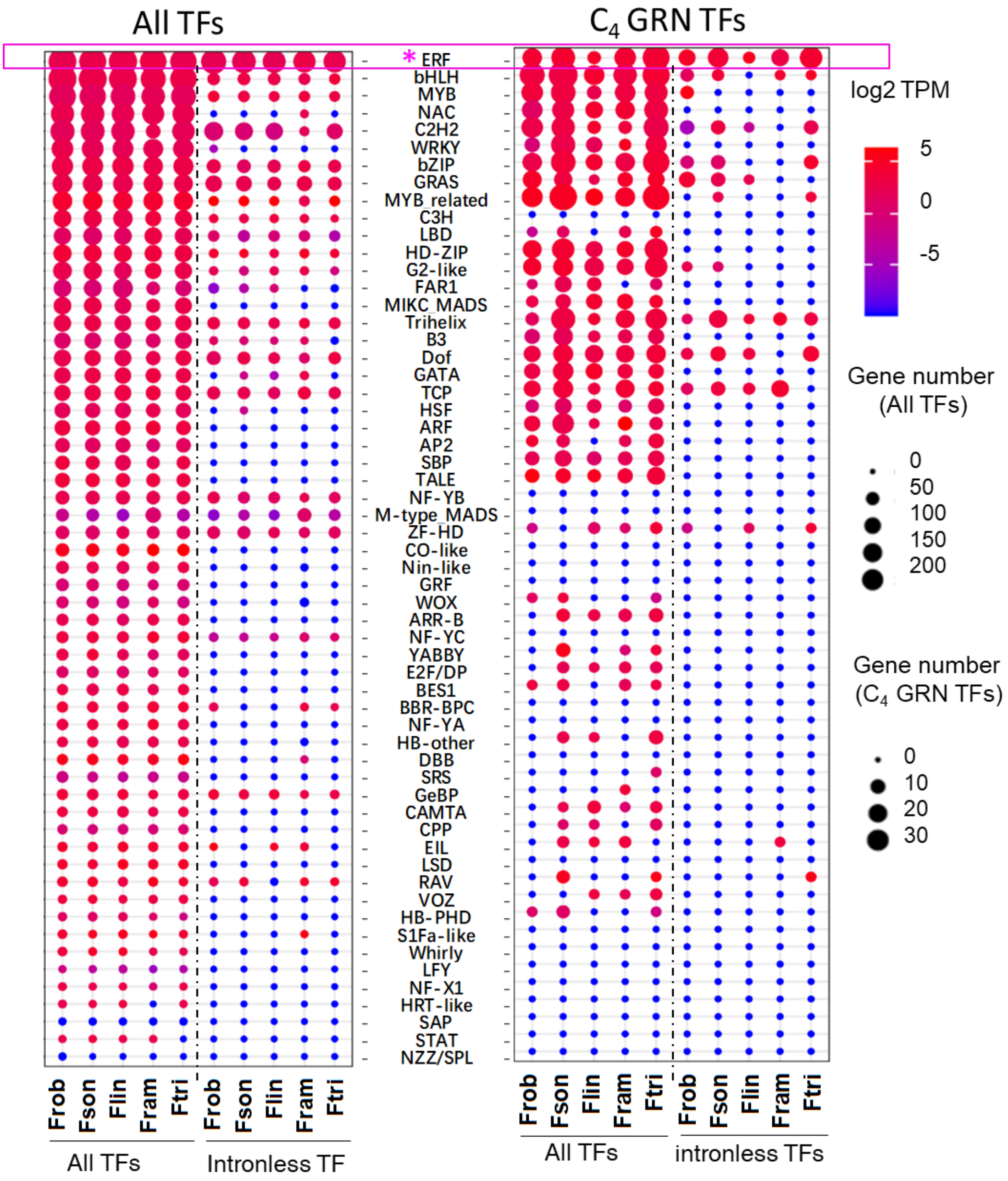


**Figure S5. The number and transcript abundances of intron-containing and intronles genes in each TF family**

Heat maps show the gene number and transcript abundance of genes in each TF family from all the annotated TFs (left panel) and from C4GRN (right panel). The size of circle represents the number of genes, and the color represents the log2 transformed transcript abundances in transcript per million mapped reads (TPM).

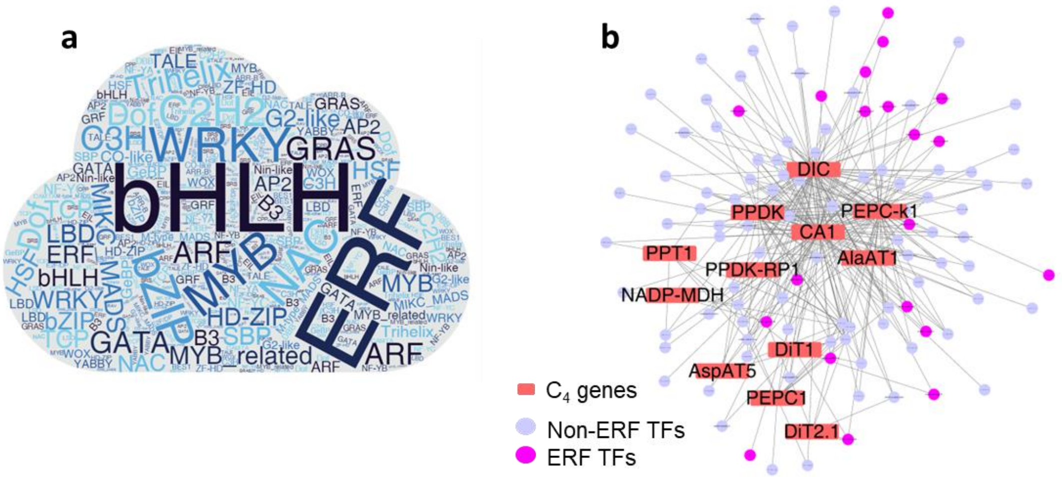


**Figure S6. ERF TF were recruited by $C_4$ genes in *Zea mays***
(a) Word cloud shows the frequencies of TFs in each TF families in the leaf gene regulatory network (GRN) of *Zea mays* (Zmay, corn) from (Tu et al., 2020). In the whole leaf GRN, bHLH is the most prevalent TFs with 138 genes. (b) The $C_4$GRN in Zmay that includes $C_4$ genes and their regulatory TFs. ERF is the most abundant TFs in the $C_4$GRN as showed in purple circle. 12 $C_4$ gene are included within the whole leaf GRN. (Abbreviation: Zmay: *Zea mays*)

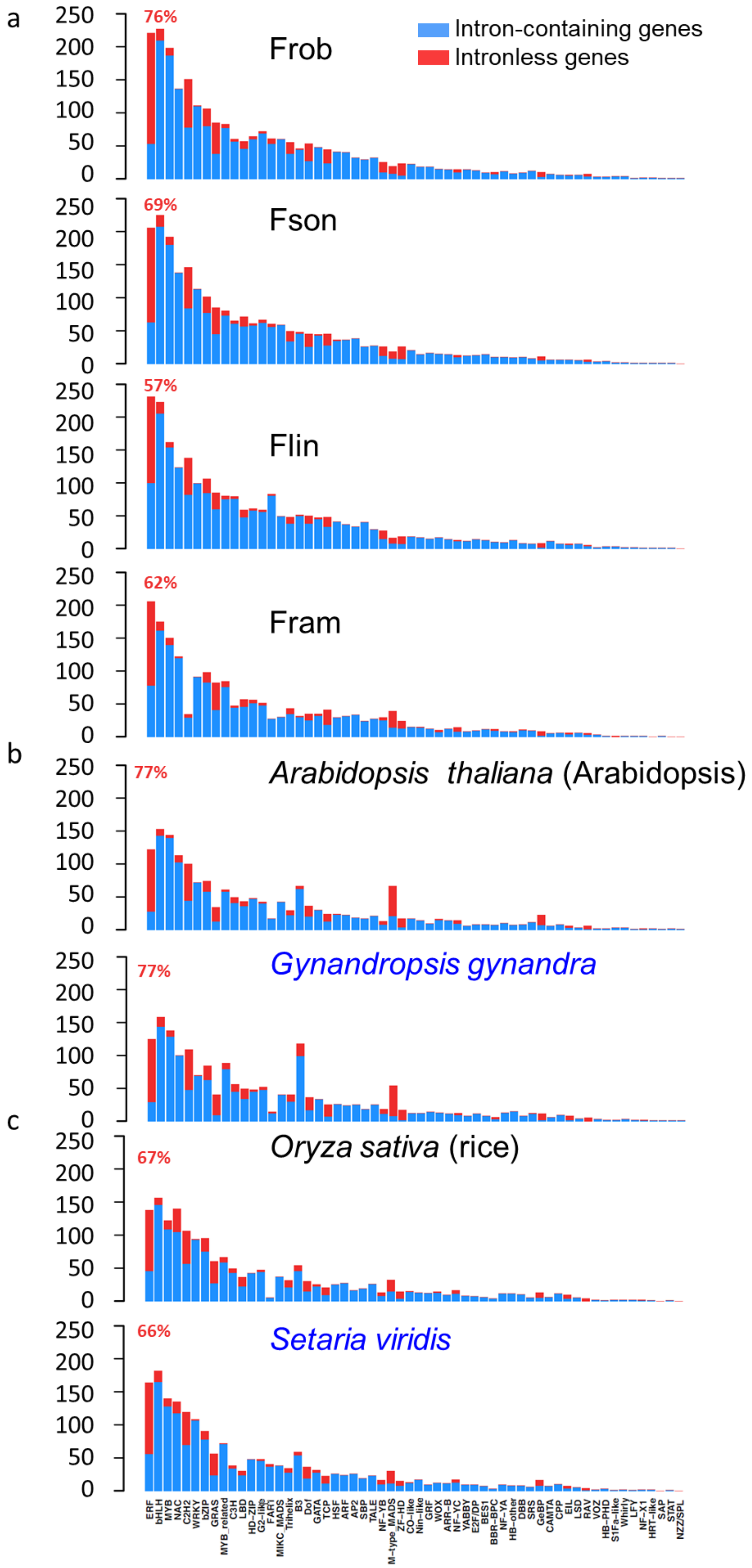
a
Frob
76%
Intron-containing genes
Intronless genes
Fson
69%
Flin
57%
Fram
62%
b
*Arabidopsis thaliana* (Arabidopsis)
77%
*Gynandropsis gynandra*
77%
c
*Oryza sativa* (rice)
67%
*Setaria viridis*
66%
250
200
150
100
50
0
ERF
bHLH
MYB
NAC
C2H2
WRKY
bZIP
GRAS
MYB_related
C3H
LBD
HD–ZIP
G2–like
FAR1
MIKC_MADS
Trihelix
B3
Dof
GATA
TCP
HSF
ARF
AP2
SBP
TALE
NF–YB
M–type_MADS
ZF–HD
CO–like
Nin–like
GRF
WOX
ARR–B
NF–YC
YABBY
E2F/DP
BES1
BBR–BPC
NF–YA
HB–other
DBB
SRS
GeBP
CAMTA
CPP
EIL
LSD
RAV
VOZ
HB–PHD
S1Fa–like
Whirly
LFY
NF–X1
HRT–like
SAP
STAT
NZZ/SPL

**Figure S7. The number of intronless genes in each TF family in different species**

The number of intronless gene and intron-contain genes are showed in each TF family from (a) four *Flaveria* species, (b) two dicotyledonous species and (c) two monocotyledonous species. $C_4$ species are labeled in blue font. Proportions of intronless genes in ERF TF family are showed in red fond for each species.

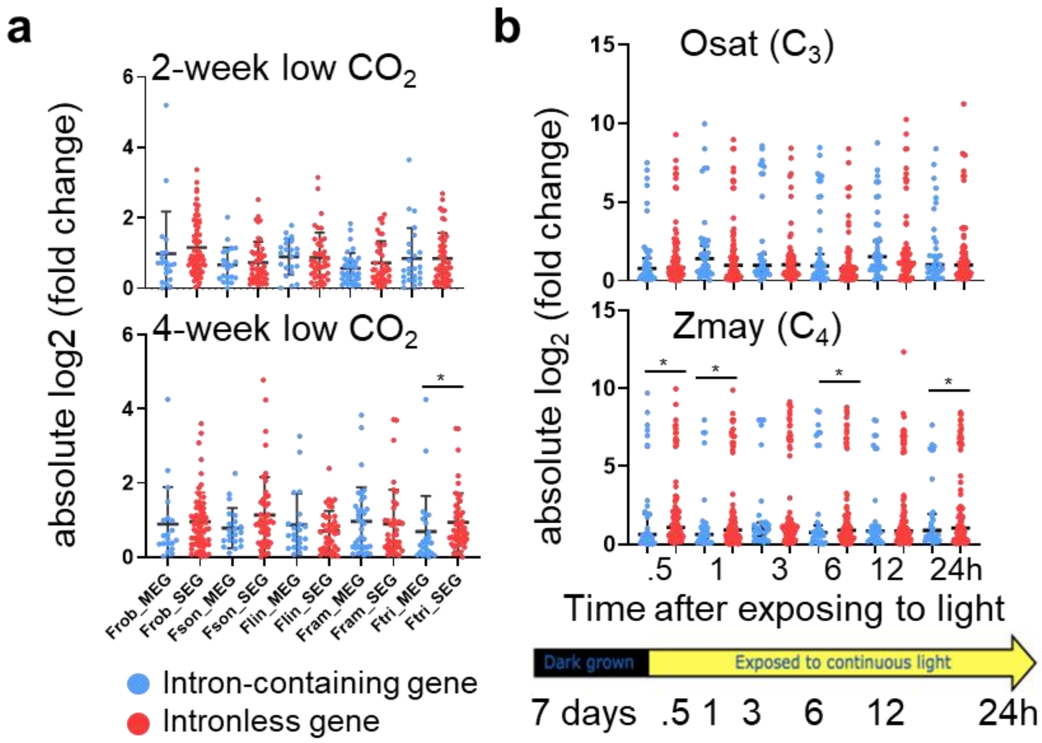


**Figure S8. The response of intronless and intron-containing ERF TFs to environmental changes**

(a) The changes on transcript abundances of intronless and intron-containing ERF TFs in five *Flaveria* species in response to low $CO_2$ (100 ppm) compared to normal $CO_2$ (380 ppm). RNA-seq data of both low $CO_2$ and normal $CO_2$ grown plants were taken from leaves after plants being grown under each condition for two weeks and four weeks respectively. (b) The response of transcript abundances of intronless and intron-containing ERF TFs in *Oryza sativa* (Osat, $C_3$) and *Zea mays* (Zmay, $C_4$) under light induction. The gene expression data of Osat and Zmay are from (Xu et al., 2016). Seeds of both species were germinated and grown under dark for 7 days. RNA-seq data of leaves were taken before light, 0.5h, 1, 3h, 6h, 12h and 24h after light respectively. The fold change of each time point was calculated as the ratio of gene expression level of this time point to that of the prior time point. Gene expression levels for all analysis was showed in transcript per million mapped reads (TPM). (Abbreviations: MEG: multi-exon genes, *i.e.*, intron-containing genes; SEG: single exon genes, *i.e.*, intronless genes; Osat: *Oryza sativa*; Zmay: *Zea mays*.)

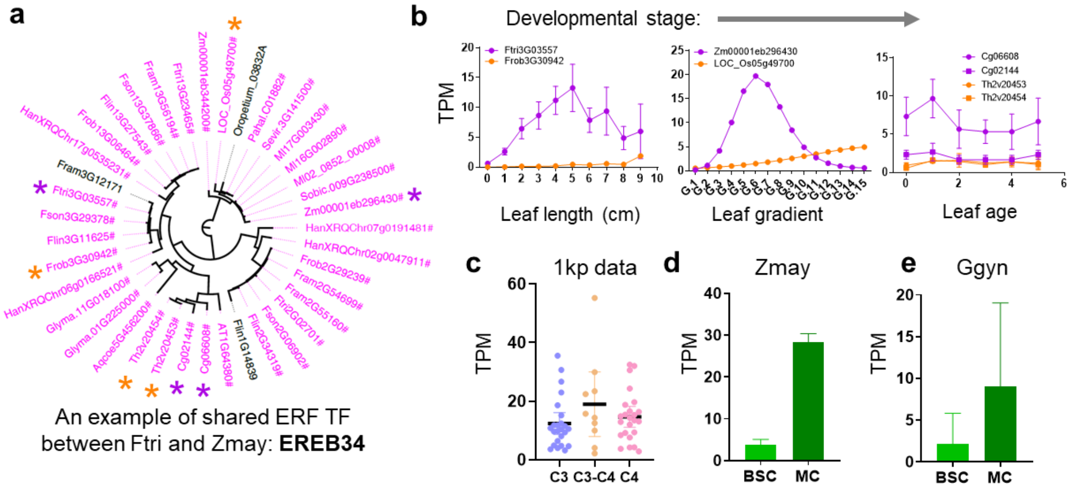


**Figure S9. An example of intronless ERF TF that was recruited to regulate $C_4$ genes in both Ftri and Zmay**

**(a)** Gene tree of EREB34. Intronless genes are labeled in purple. EREB34 orthologous genes among 23 species (see Figure5) were predicted applying Orthofiner. EREB34 present in higher plants but not in algae or liverwort. Genes marked with orange and purple stars are from $C_3$ and $C_4$ plants that compared in transcript abundances in **(b)**. **(b)** Comparisons of EREB34 in transcript abundances between Frob ($C_3$) *vs* Ftri ($C_4$) (lef), *Oryza sative* (Osat, $C_3$) *vs Zea mays* (Zmay, $C_4$) (middle), and *Tarenaya hassleriana* (Thas, $C_3$) vs *Gynandropsis gynandra* (Ggyn, $C_4$) (right) along leaf developmental gradient. Genes from $C_3$ species are labeled in orange, and those from $C_4$ species are in purple. RNA-seq data of these species are from published sources, *i.e.*, data of Frob and Ftri are from (Billakurthi. et al., 2020), data of Osat and Zmay are from (Xu et al., 2016), data of Thas and Ggyn are from (Kulahoglu et al., 2014) . The first points in Frob and Ftri (0) represent meristem. Leaf ages of Thas and Ggyn are as following: **0**: 0-2 days (d); **1**: 2-4 d; **2**: 4-6 d; **3**: 6-8 d; **4**: 8-10 d and **5**: 10-12 d. **(c)** Transcript abundances of EREB34 in $C_3$, $C_3$-$C_4$ and $C_4$ species from one thousand plants (1kp) project, covering 18 independent $C_4$ lineages. RNA-seq data are from (Steven Kelly, 2018). **(d)** Transcript abundance of EREB34 in Zmay bundle sheath cell (BSC) and mesophyll cell (MC). Expressional data are from (Chang et al., 2012). **(e)** Transcript abundances of EREB34 in Ggyn BSC and MC. Expressional data are from (Aubry et al., 2014). (Abbreviations: MC: mesophyll cell; BSC: bundle sheath cell.)
